# After the honeymoon, the divorce: unexpected outcomes of disease control measures against endemic infections

**DOI:** 10.1101/608653

**Authors:** Brandon Hollingsworth, Kenichi W Okamoto, Alun L Lloyd

## Abstract

The lack of effective vaccines for many endemic diseases often forces policymakers to enact control programs that rely on non-immunizing controls, such as vector control, in order to reduce the massive burden of these diseases. It is well known that controls can have counterintuitive effects, such as the honeymoon effect, in which partially effective controls cause not only a greater initial reduction in infection than expected for an infection near its endemic equilibrium, but also large outbreaks during control as a result of accumulation of susceptibles. Unfortunately, many control measures cannot be maintained indefinitely, and the results of cessation are not well understood. Here, we examine the results of stopped or failed non-immunizing control measures in endemic settings. By using a mathematical model to compare the cumulative number of cases expected with and without the control measures, we show that deployment of control can lead to a larger total number of infections, *counting from the time that control started*, than without any control – the *divorce effect*. This result is directly related to the population-level loss of immunity resulting from non-immunizing controls and is seen in model results from a number of settings when non-immunizing controls are used against an infection that confers immunity. Finally, we also examine three control plans for minimizing the magnitude of the divorce effect in seasonal infections and show that they are incapable of eliminating the divorce effect. While we do not suggest stopping control programs that rely on non-immunizing controls, our results strongly argue that the accumulation of susceptibility should be considered before deploying such controls against endemic infections when indefinite use of the control is unlikely. We highlight that our results are particularly germane to endemic mosquito-borne infections, such as dengue virus, both for routine management involving vector control and for field trials of novel control approaches.

**Author Summary:** Many common endemic infections lack effective, inexpensive vaccinations, and control relies instead on transmission reduction, e.g. mosquito population reduction for dengue. Often, these controls are used with the immediate goal of decreasing the current incidence with little importance placed on what will happen at later points in time, and much less what will happen once the control is stopped. Here, by looking at the cumulative incidence since the beginning of the control period, instead of the instantaneous incidence, we show that when controls are stopped, or fail, the resulting outbreaks can be large enough to completely eliminate any benefit of the control. We call this result the *divorce effect*. Further, we show that this result is not limited to specific transmission pathways or epidemiological parameters, but is instead tied directly to the reduction of herd immunity inherent in non-immunizing controls. Lastly, by evaluating programs to minimize the magnitude of the divorce effect, we show that without maintaining herd immunity, or successfully continuing control for decades, it is impossible to keep the costs of post-control outbreaks from outweighing the benefits of the control program.

## Introduction

An estimated 200 million cases of malaria, 390 million cases of dengue fever, and 9 million cases of measles occurred in 2016 [1,2], representing only a portion of the total impact of endemic disease that year. The burden that this places on local populations, both in terms of morbidity and mortality and both direct and indirect economic costs, often pressures policy makers to act to suppress these infections. However, the scientific rationale on which the implemented policies are based is not always clear, making it difficult to assess whether the risks associated with control have been adequately addressed.

Eradication— the permanent reduction of worldwide incidence to zero [3]— is the ideal aim of all control programs. This goal is unrealistic, with only two infections having been successfully eradicated to date: smallpox and rinderpest [4]. Often, a more realistic goal for a control program is either long-term suppression or local elimination of the infection. These goals hold their own challenges though, as they require long-term or even indefinite control programs, which can face budgetary and public support issues, not to mention the potential for some controls to fail due to evolution of resistance. Further, if there is a loss of herd immunity in the population due to the control lowering population exposure to the pathogen, there is the additional risk that when a control program ends the infection will re-emerge in a post-control epidemic and reestablish in the population [4].

Naively, one might imagine that lowering the incidence of infection will have no detrimental effects for the population. However, mathematical modeling has previously revealed numerous perverse outcomes of application of ineffective control measures (by which we mean ones that do not bring the basic reproductive number, *R*_0_, below one) in endemic settings. Perhaps the most famous example is the increased age at infection that results when a population is partially vaccinated for rubella, leading to more infections occurring in women of child-bearing age, where severe complications, such as congenital rubella syndrome, can result when pregnant women become infected [5–7]. While this certainly represents a potential downside of the control, the population sees a reduction in rubella prevalence. McLean and Anderson (1988) showed that when an ineffective control is used against an endemic infection it often results in an initial drop in prevalence to well below the endemic level, the “honeymoon effect”, but this is followed by outbreaks that periodically increase prevalence above the endemic level as a consequence of a build-up of susceptible individuals. Similarly, in a seasonally-forced setting, Pandey and Medlock [9] found that vaccination against dengue virus could result in a transient period with periodic outbreaks of larger peak prevalence than occurred before vaccination. These last two examples illustrate possible negative side effects of ineffective controls: they can cause transient increases in prevalence while still resulting in a decrease in total incidence.

In the results above, there is higher incidence than expected, but Okamoto et al.[10] described an even more troubling theoretical result while exploring a model of failed or stopped combined strategies aimed at controlling dengue virus, e.g. vaccination along with transgenic vector control. They observed that when control was only transient the **total** number of infections that occurred, counting from the time that control started, a quantity they called the cumulative incidence (CI), could exceed the number of cases that would have been observed had no control been deployed. Even in situations where control measures had a significant positive impact over a period of years, the outbreaks that ensued following the cessation, or failure, of control could lead to an outbreak that was large enough to outweigh the number of cases prevented during the control period.

While Okamoto et al. [10] showed that it was possible for transient transgenic controls to increase the total number of infections, here we demonstrate that this effect—which we call the *divorce effect*—is not an artifact of very specific complex models, but quite a general phenomenon that can occur across a range of models and parameter space when deploying a control measure that does not confer immunity. By exploring the dynamics of the divorce effect in the setting of several simple models we gain insights that were not obtainable using the previous complex models. Conversely, we find that for immunizing controls (e.g. vaccination) the divorce effect does not occur, even when the duration of protection is relatively short-lived.

We demonstrate the generality of this result for endemic infections by simulating cessation of control measures in three commonly-used models for pathogen transmission. Unlike the honeymoon effect, the divorce effect occurs for both ineffective and effective controls, provided that they are transient. As anticipated, control results in the accumulation of susceptible individuals resulting in the potential for a large outbreak following the cessation of control. This outbreak is either triggered by infective individuals that remain in the population or by reintroduction of infection from outside the control area, and its size increases asymptotically towards the size of a virgin-soil epidemic as the length of the control period is increased and herd immunity is lost. Counterintuitively, and comparable to results in Okamoto et al. [10], we see that the post-control outbreak often results in there being timeframes over which the cumulative incidence of infection since the start of control is higher than would have occurred in the absence of control. Further, these outbreaks are significantly larger than the endemic levels of the infection and would likely overwhelm healthcare providers in the area.

This paper is organized as follows. We first describe the three models we choose to illustrate the divorce effect: a non-seasonal SIR model, a seasonal SIR model, and a host-vector model. We then demonstrate, in each setting, the occurrence of the divorce effect and its sensitivity to relevant parameters, namely *R*_0_ and the duration and strength of control. Further, for the seasonal SIR model, we explore the sensitivity of the strength of the divorce effect on the timing of the start and end of the control. Then for the seasonal SIR and seasonal host-vector model we look at three possible strategies for mitigating the divorce effect and show they are incapable of eliminating the divorce effect. A crude analytical approximation for the divorce effect and additional models are explored in the Supplemental Information, as is the impact of using immunizing controls.

## Models

To evaluate the magnitude of the Divorce Effect, we simulate the cessation of a short-term control affecting transmission in three infection systems: a SIR model, a seasonal SIR model, and a host-vector SIR model. While these are the only models we discuss in detail here, this result can be seen in most models that have a replenishment of the susceptible population, including the more general SIRS model, for which host immunity is not life-long, and an age-structured model with realistic mixing parameters (see Supplemental Information for exploration of additional forms of transmission models). These results are parameterized for a human population and mosquito vector, but the results are generalizable to other species.

### SIR Model

We assume a well-mixed population of one million hosts and a non-fatal infection that is directly transmitted and confers complete life-long immunity. The numbers of susceptible, infective, and removed individuals are written as *S*, *I* and *R*, respectively. We allow for replenishment of the susceptible population by births, but assume the population size is constant by taking per-capita birth and death rates, μ, to be equal (this assumption is relaxed in the supplemental information). This results in the standard two-dimensional representation of the SIR model, where the number of removed individuals is *R* = *N* − *S* − *I* (Equation 1).

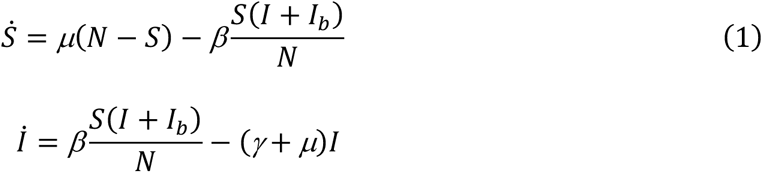

For our simulations, we assume parameters resembling a short-lived infection in a human population, lasting on average 5 days (average recovery rate, γ = 73/year) and that individuals live on average 60 years (µ = .0167/year), allowing the transmission parameter, *β*, to be adjusted to achieve the desired value of *R*_0_. In order to reseed infection following cessation of control and to counter the well-known weakness of infective numbers falling to arbitrarily low levels in deterministic transmission models, we follow numerous authors in including a constant background force of infection [11,12] in the model. This represents infectious contacts made with other populations, and occurs at a rate that is equivalent to there being *I*_b_ additional infective individuals within our focal population. For our simulations, we take *I*_*b*_ = 1(sensitivity of our results to *I*_*b*_ can be found in the supplemental information).

### Seasonal SIR Model

For the seasonal SIR model, we allow the transmission parameter to fluctuate seasonally (annually) around its mean, *β*_0_, taking the form given in Equation 2. Seasonal oscillations in the parameter have relative amplitude *β*_1_ = .02 with maxima occurring at integer multiples of 365 days. Noting that seasonally forced models are particularly susceptible to having the number of infectives fall to unreasonably low numbers between outbreaks [13], we again take *I*_*b*_ = 2 in the background force of infection term.

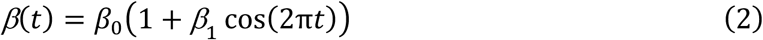

### Host-Vector Model

We model an infection with obligate vector transmission. As in other models, we assume that the host population size is held constant (*R* = *N* − *S* − *I*), but we allow the vector population size to fluctuate—so that, for instance, we can model vector control. For simplicity, we only model the female adult vector population and assume density-dependent recruitment into the susceptible class (*U*), with a logistic-type dependence on the total female adult population size. Infectious vectors (*V*) arise from interactions with infected hosts (Equation 3).

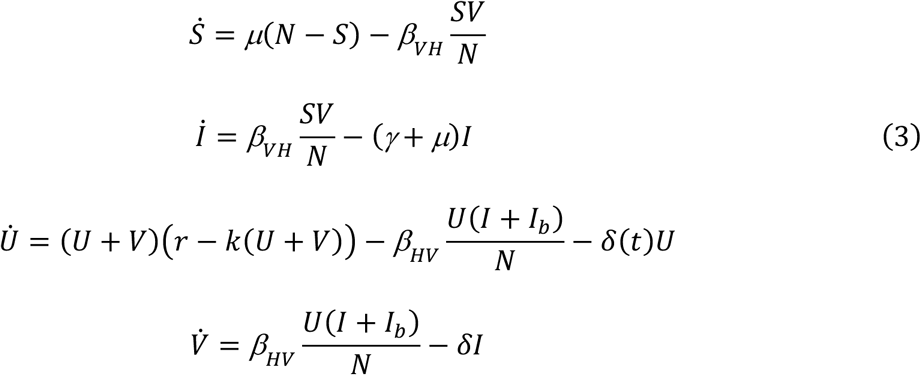

We assume that host demography and recovery rates are the same as in the SIR model, with a host population of one million individuals. We assume that the vector lives on average 10 days (*δ* = 36.5/year), the growth constant (*r*) and density dependence parameter (*k*) are parameterized as in Okamoto et al. (2016): *r* = 304.775/year and *k* = 1.341×10^−7^/(vector*year), resulting in an equilibrium vector population of 2 million individuals. The transmission parameter from host to vector (*β*_*HV*_) is assumed to be 109.5/year and the parameter for vector to host (*β*_*VH*_) is changed to produce the desired *R*_0_. We again assume a background force of infection (with *I*_*b*_ = 2), representing reintroduction of infection from outside our focal population.

Seasonality plays a large role in vector-borne infections and affects many aspects of the infection and its vector. Temperature affects breeding rates, larval development, and death rates of the vector, the extrinsic incubation period and transmissibility of the infection itself, and host encounter rates, while precipitation can affect the availability of appropriate habitat and encounter rates [14–16]. However, most of these add a level of model complexity which is unnecessary for this study, so we choose to use a simple forcing term for mosquito recruitment that fluctuates seasonally with relative magnitude *r*_s_ (*r*_*s*_ = 0.02) about its baseline (*r*_0_ > = 304.775 /year) (Equation 4).

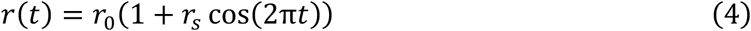

### Control

We model a control that is applied instantaneously and consistently from time *t*_0_ (which, for simplicity, we usually take to be equal to zero) to time *t*_end_ and is instantaneously removed at the end of the control period. In the SIR and seasonal SIR models, control reduces the transmission rate by some proportion, *ε*, and, in the host vector model, causes a proportional increase, *σ*, in the vector mortality rate. This results in the transmission parameter given in Equation 5 for directly transmitted infections and the vector death rate given in Equation 6 for the vector-borne infections.

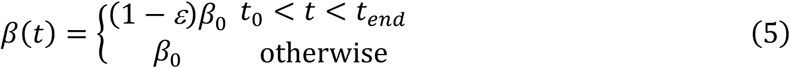

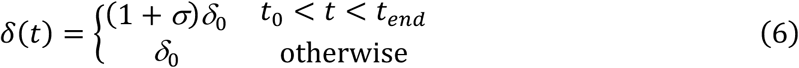

While we only look at these control measures in the main text, other controls (such as an increase in the recovery rate, *γ*) are explored in the Supplemental Information (Figure S2), and give similar results.

### Measuring Effectiveness

There are a number of measures that can be used to quantify the effectiveness of a control. We want to characterize the total number of cases that occur from the start of control until a particular point in time, a quantity we call the cumulative incidence (CI). For a directly transmitted infection, this is calculated as follows

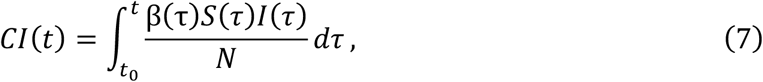

i.e. by integrating the transmission term over the time interval from the start of control until the time, *t*, of interest. This quantity could be calculated both in the presence of control and in the baseline, no-control, setting; we distinguish between these two by labeling quantities (e.g. state variables) in the latter case with a subscript B to denote baseline.

One commonly-used measure of effectiveness is the number of cases averted by control (CA), *CI*_B_(*t*) - *CI*(*t*). This has the disadvantage (particularly in terms of graphical depiction) that it can become arbitrarily large as *t* increases. Consequently, some authors choose to utilize a relative measure of cases averted, dividing by the baseline cumulative incidence (see, for instance, the work of Hladish et al. [17]). We instead follow our earlier work and use the relative cumulative incidence (RCI) measure employed by Okamoto *et al.* [10], calculating the cumulative incidence of the model with the control program relative to the cumulative incidence of the model without the control program (Equation 8).

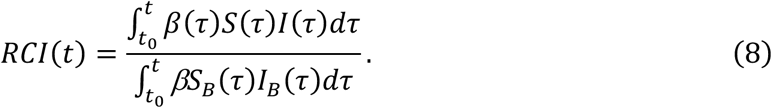

RCI(*t*) values above one imply that the control measure has resulted in an increase in the total number of cases compared to the baseline. Importantly, as time becomes larger, RCI becomes less sensitive to outbreaks in the system. For a transient control, RCI will approach 1 as *t* becomes larger.

We see that the relative cases averted measure employed by Hladish et al. [15] is simply 1-RCI(*t*). Both relative measures have properties that make them attractive for graphical depiction although it should be borne in mind that both involve a loss of information on the actual number of cases averted. For example, an RCI of 1.1 after one year is a much smaller increase in total cases than an RCI of 1.1 after 10 years, and an RCI of just below one after many years can represent a large reduction in total incidence. In cases where this information is pertinent, it may be more appropriate to use non-relative measures such as cases averted. The choice of measure does not impact the occurrence of the divorce effect; figures that show cases averted are included in the Supplemental Information (Figure S1).

Analogous expressions for CI and RCI can be written for the host-vector model using the appropriate transmission terms.

## Results

### SIR Model

Simulations show the successful suppression of infection following the implementation of a control which reduces the transmission parameter, *β*, in the population. With infection at endemic equilibrium, the honeymoon effect [8] states that even a modest reduction in the transmission parameter will have a large effect on the incidence of the infection due to the effective reproductive number, *R*_*t*_, the expected number of new infections each infectious individual causes, being one. After the control is stopped, the incidence of the infection remains low for some time as the number of infective individuals builds from very low numbers (Figure 1(a), curve). However, once control ends *R*_*t*_ immediately rises above one and continues to increase while prevalence is low, due to the buildup of the susceptible population (plots of *R*_*t*_ and *S*(*t*) are provided in the supplemental information: see Figure S3). This increased *R*_*t*_ eventually drives a large outbreak, quickly depleting the susceptible population, at which point incidence (Figure 1(a), black curve), and *R*_*t*_, again fall to low numbers.

**Figure 1:**
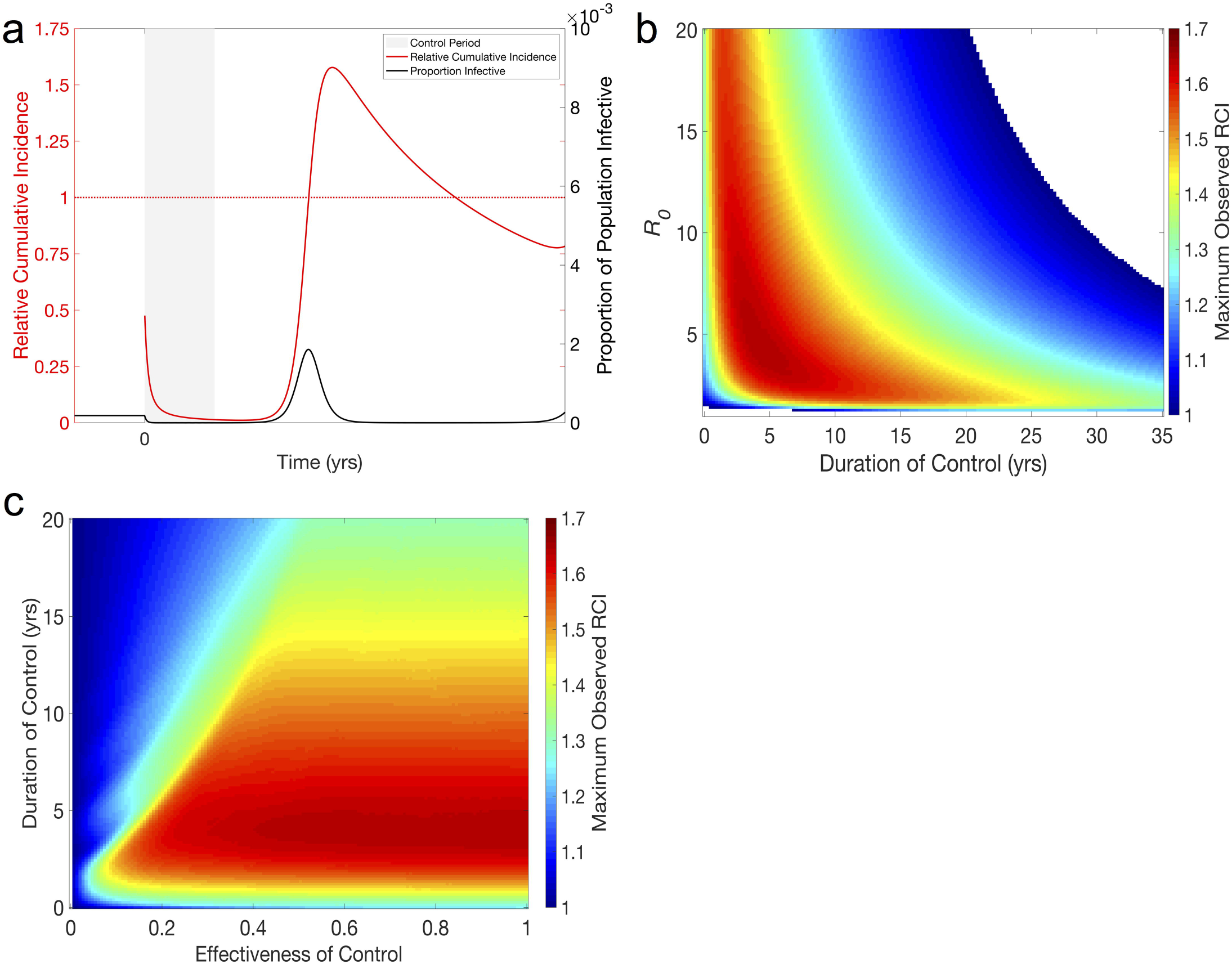
The divorce effect in the SIR model. (a) Typical time-series showing the divorce effect. Beginning at time zero, a year-long 50% reduction in the transmission parameter of an endemic infection (*R*_0_ = 5, *β* = 365 /year, *γ* = 73 /year) reduces prevalence of the infection to near zero for the length of the control, where it remains until time 1.5 yrs, at which point a large post-control outbreak occurs. RCI falls towards zero as prevalence remains low, but the post-control outbreak is large enough to bring RCI well above 1 (peak RCI is approx .1.4). **(b) Magnitude of divorce effect in terms of relative cumulative incidence (RCI).** Maximum RCI is found as the highest value of RCI observed within 25 yrs following a 100% effective control of an infection with 1<*R*_0_<20 and lasting between 1 month and 35 years. RCI>1 indicates the divorce effect and we see that the divorce effect occurs across a large portion of the parameter space, and ubiquitously for controls lasting less than 20 years. *β* is varied to attain the desired *R*_0_, all other parameters as in (a). **(c) Maximum RCI for a given effectiveness and duration of control.** The maximum RCI is found as the maximum observed RCI within 25 yrs after the end of a control that is between 0% and 100% effective and lasts between 1 month and 20 years (*R*_0_=5). The ridge between areas of high and low maximum RCI results from ineffective controls being maintained long enough for outbreaks due to the honeymoon effect deplenishing the population of susceptible individuals before the control periods end. All other parameters as in (a).

To evaluate the success of the control, we examine the RCI in the period following introduction of control and see that during and immediately following the control period, when incidence is low, the RCI decreases towards 0, suggesting a successful control program. However, once the post-control outbreak begins, RCI increases rapidly resulting in the divorce effect (RCI>1) before dropping back below one once the epidemic begins to wane and incidence falls below endemic levels (Figure 1(a)). During the period where RCI>1, lasting approximately 2 years in our example, the control has not only failed to decrease the total incidence of infection but has resulted in an increase in total incidence, the divorce effect. Following this initial outbreak and trough, RCI continues to oscillate around one, and approaches one in the long run (see Figure S4).

Exploring values of *R*_0_ and the duration and strength of control shows that the divorce effect is present over a wide region of parameter space. Figure 1(b) shows the magnitude of the divorce effect, quantified by the maximum RCI seen, as a function of *R*_0_ and duration of control for a perfect control measure (*β* = 0 during the control period). Perfect control was employed here to eliminate any confounding effects from the honeymoon effect that could occur during an imperfect control. We find that for the most biologically relevant area of parameter space (*R*_0_<20, control lasting less than 20 yrs) the divorce effect always occurs and will result in a 20-60% increase in cumulative incidence (RCI=1.2-1.6) at its peak. However, we also find that it is possible to avoid the divorce effect if controls are maintained long enough. For infections with a high *R*_0_, this requires maintaining the control for decades, and the length of time needed grows as *R*_0_ is decreased. The non-monotonic relationship between the magnitude of the divorce effect and the length of the control seen here suggests that a control program should either be discontinued immediately, if *R*_0_ is small, or continued as long as possible to avoid the divorce effect (Figure 1(b); see also Figure S5 in Supplemental Information).

Relaxing our assumption of a completely effective control and focusing on a fixed *R*_0_ (*R*_0_=5, Figure 1(c)), we see that the relationship between the magnitude of the divorce effect and the length of the control period varies with the strength of the control. A steep edge-like pattern is seen in Figure 1c when control is ineffective but carried out for a long period of time, a consequence of the honeymoon effect. For populations at endemic equilibrium, the honeymoon effect means that any reduction in transmission will be sufficient to significantly reduce transmission for a period of time. For controls that are relatively short lived, here approximately 5 years, the control does not outlast the honeymoon period, resulting in the magnitude of the divorce effect being relatively insensitive to the effectiveness of the control in this region of parameter space. How the interaction between the effectiveness of control and *R*_0_ affects the magnitude of the divorce effect is explored in the supplemental information (Figure S6).

### Seasonal SIR Model

Temporary control measures in the seasonal SIR model show many of the same dynamics as in the non-seasonal model, namely that a successful control is followed by a period of low incidence and eventually a post-control outbreak leading to a divorce effect (Figure 2(a)) before settling back into regular seasonal outbreaks (Figure S7). However, the timing and size of the post-control epidemic, and thus the magnitude of the divorce effect, depend not only on *R*_0_ and the length of the control but also the timing of both the onset and end of the control (Figures 2(b) and 2(c)). This leads to a highly nonlinear dependence of the magnitude of the divorce effect on *R*_0_ and the duration of control (Figure 2(b)). However, the presence of ranges of parameter space with smaller magnitudes of the Divorce Effect at regular intervals could allow policy makers to determine optimal times to stop control. These effects become more apparent with an increase in seasonality (Figure S8). As seasonality increases, the differences due to timing become more pronounced, resulting in more potential for mitigating the divorce effect with a properly timed treatment. Conversely, this also means a larger divorce effect will be seen with a poorly timed treatment (Figure S8).

**Figure 2:**
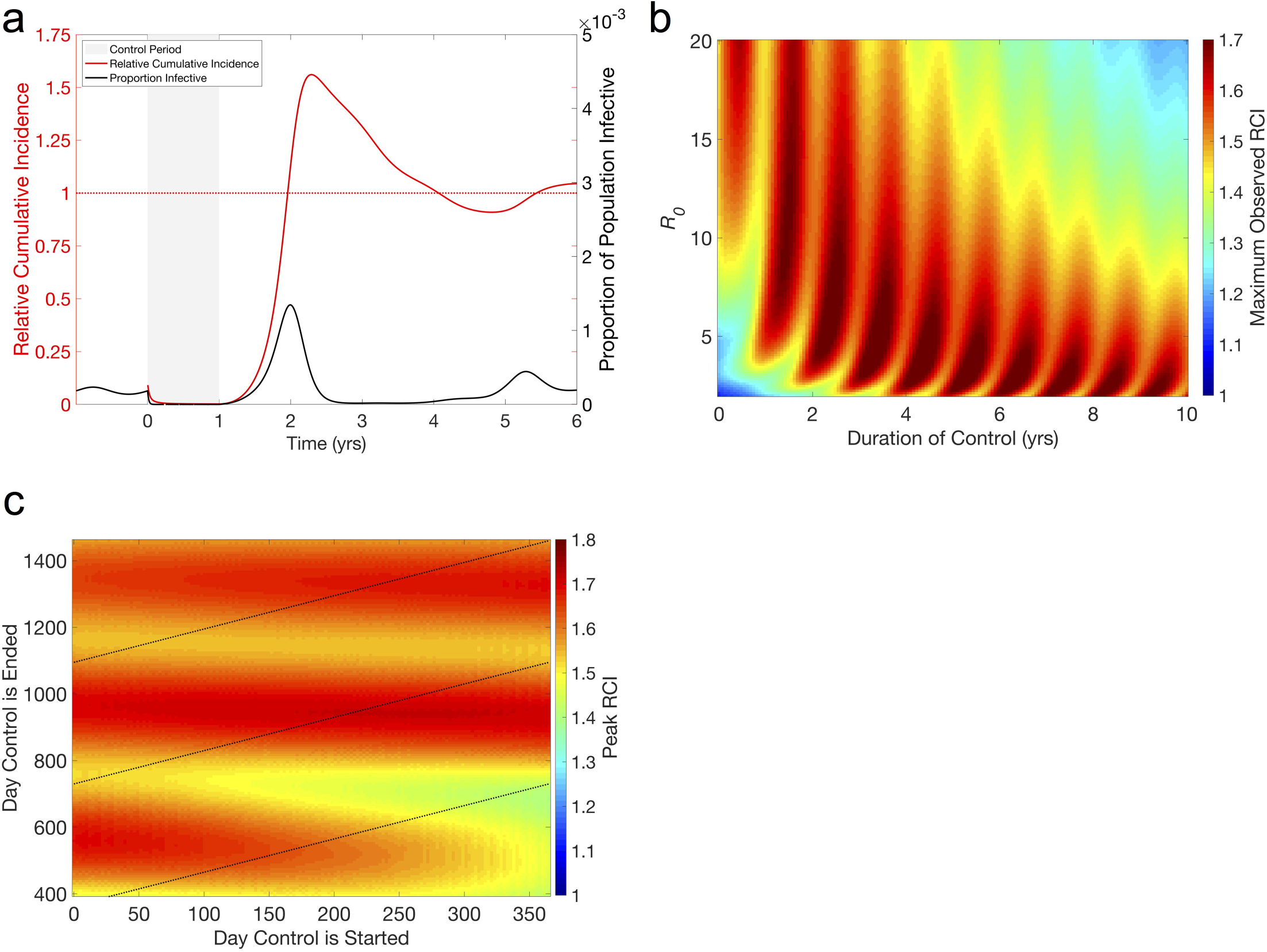
The divorce effect in the seasonal SIR model. (a) Typical time-series showing the divorce effect. Beginning at time zero, when the transmission parameter is at its maximum, a year-long 90% reduction in the transmission parameter of an endemic infection (*R*_0_ = 5, β_0_ = 365 /year, β_1_ = .02, γ = 73/year) is implemented at the beginning of a seasonal outbreak and reduces prevalence of the infection to near zero for the length of the control. Following the end of the control, a large outbreak, many times the size of the regular seasonal outbreaks, occurs during the next season. RCI falls towards zero as prevalence remains low while the control is in effect and rises above 1 during the large outbreak the following year (Maximum RCI = 1.2). **(b) Magnitude of divorce effect in terms of relative cumulative incidence (RCI).** Maximum RCI is found as the highest value of RCI observed within 25 yrs following a 100% effective control of an infection with 1<*R*_0_<20 and lasting between 1 month and 35 years. RCI>1 indicates the divorce effect and we see that the divorce effect occurs in most of the parameter space. β_>_ is varied to attain the desired *R*_0_, with all other parameters as in (a). **(c) Effect of timing on the magnitude of the divorce effect.** Maximum RCI is the highest RCI observed within 25 yrs following a 100% effective control of an infection with *R*_0_ = 10 (*β* = 730 /year, all other parameters as in (a)) beginning and ending on specified days. Dashed lines represent controls lasting either 1, 2, or 3 years. Unlike the non-seasonal SIR model (Figure 1), the magnitude of the divorce effect is not solely dependent on *R*_0_ and the length of the control. Maximum RCI is most sensitive to the day the control is ended, moderately sensitive to the day it is started, and only slightly sensitive to the length of the control. This is due to the timing of the end of the control determining the timing of the outbreak. We also see that continuing the control for another year often has little impact on the magnitude of the divorce effect.

The oscillatory nature of the relationship between the maximum RCI and *R*_0_ (Figure 2(b)) implies a relationship between the timing of the control period and the severity of the divorce effect. While the magnitude is only highly sensitive to the start time for very short control periods, lasting around a year, it is highly sensitive to the end time (Figure 2(c)). This means that controls of similar lengths can have significantly different outcomes depending on their timing, e.g. a 1 year control ending day 700 results in a maximum RCI around 1.4 while a control of the same length ending day 515 results in a maximum RCI near 1.7. This is a direct result of the seasonal forcing function and delaying the outbreak until a period in which *R*_0_ is larger, similar to results seen when controls are used against epidemics in seasonal settings [18,19]. Regardless of start time, the optimal end time occurs shortly after the peak in the transmission parameter, β(*t*), (days 750 and 1155 in Figure 2(c)), suggesting this would be the best time to end control programs.

### Host-Vector Model

The non-seasonal host-vector model has broadly similar dynamics to the non-seasonal SIR model in terms of the divorce effect (Figures S10 and S11), so here we focus instead on the seasonal host-vector model. Following one year of insecticide treatment that reduces the average mosquito lifespan by a half (i.e. increases the mosquito death rate by 100%, σ = 1) the infection is suppressed and there is no seasonal outbreak for the next two years (Figure 3). A major outbreak, with approximately eight times the peak prevalence of the pre-control seasonal outbreaks, occurs in the third year and results in a maximum RCI of around 1.50, before the epidemic fades and incidence again returns to low levels. The size of this outbreak would almost certainly risk overwhelming even the most well-funded medical services. RCI then remains above 1 until year 7. The population continues to see large periodic outbreaks, each bringing RCI back above 1, for decades until the endemic equilibrium is reached again (Figure S12).

**Figure 3:**
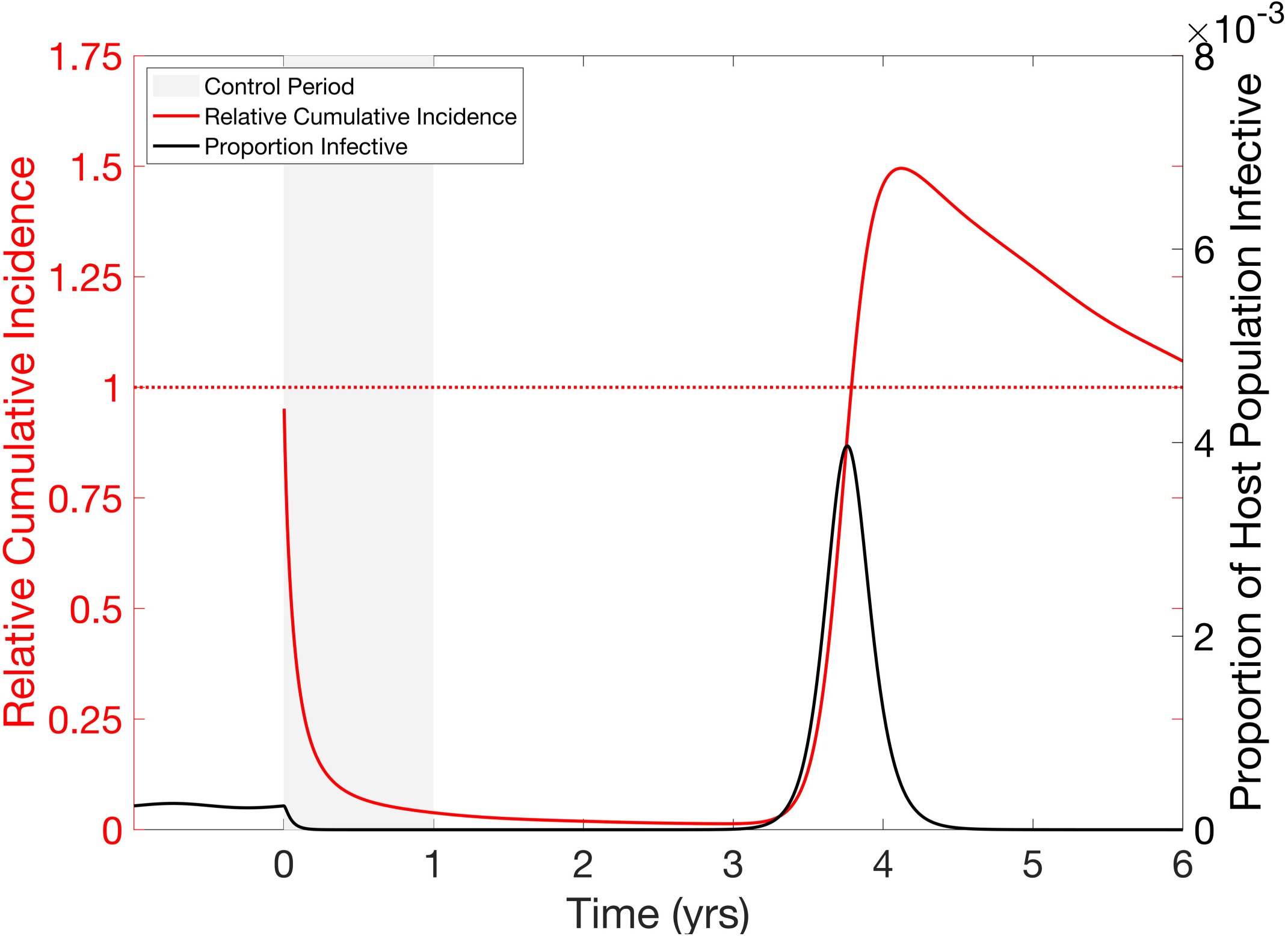
Divorce Effect in a Seasonal Host-Vector model. Control is shown in a seasonal (*r*_*s*_ = .02) host-vector model with *R*_0_ = 5. Beginning at time zero, a control is implemented that increases the vector mortality rate by 100% (corresponding to a 50% drop in vector life expectancy). This results in a reduction in prevalence (black curve) of the infection to near zero during the control period, where it remains until roughly time 3 yrs, at which point a large post-control outbreak occurs. RCI (red curve) falls towards zero during the control period and while prevalence remains low, but the post-control outbreak is large enough to bring RCI above 1 (peak of approx. 1.49).

### Mitigating the Divorce Effect

It is apparent from earlier results (e.g. Figure 1(b)) that avoiding the divorce effect in a non-seasonal setting is only possible with a non-immunizing control by maintaining suppression for decades, due to the inevitable build-up of susceptible individuals. Therefore, the goal in these situations should be to maintain the control as long as possible or until a vaccine becomes available, and we focus instead on the seasonal SIR and host-vector models. In this section, we look at three different treatment plans for deploying a set amount of treatment, twelve one-month treatments, and their ability to mitigate the divorce effect. The first relies on annual controls lasting one month when *R*_0_ is at its maximum, the second has a month-long control applied in response to the prevalence reaching some set level—which we might imagine corresponding to an outbreak becoming detectable or reaching a sufficient level to cause concern to local authorities—that we take here to be when two hundred individuals out of a million are infective, and the third chooses when to implement a month-long control based on minimizing the peak RCI. For comparison, all three use 12 total months of control.

With annual monthly control for a directly transmitted seasonal infection, the population sees a significant initial reduction in prevalence. However, as predicted by the honeymoon effect, the repeated use of controls results in a diminished effect on the prevalence and seasonal outbreaks begin to occur between control periods. The peak prevalence of these outbreaks quickly grows to be significantly larger than the seasonal outbreaks before the control program was begun, however they are blunted by the next control period before RCI rises above one. Once the program is ended, however, a post-control outbreak quickly brings RCI above one (Figure 4(a)).

**Figure 4:**
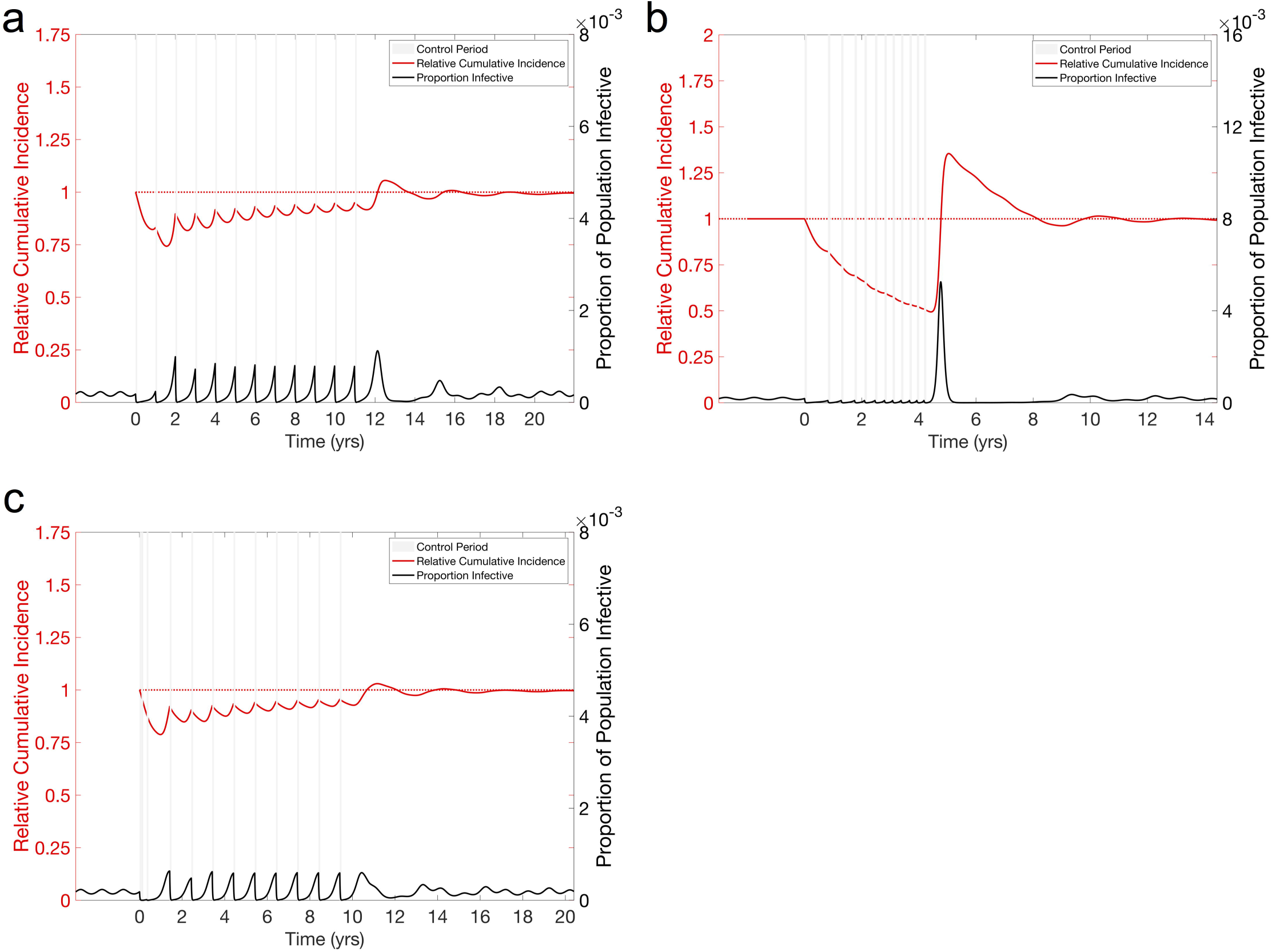
Suggested techniques for mitigating the divorce effect with seasonal transmission. We consider an endemic disease, parameterized as in Figure 2(a). In all cases, twelve 1/12 yr. controls are used, to be consistent with the 1 yr. controls used in other figures, reducing the transmission parameter by 90% (ϵ = .9). **(a) Pulsed control for Seasonal SIR model.** Control occurs yearly at a fixed time (when *R*_0_ is highest) for a fixed time (1/12 yr.) to control an endemic disease (parameterized as in Figure 2(a)). The control is effective at stopping the outbreak the first year, but seasonal outbreaks in subsequent years are larger, driven by an increasing population of susceptible individuals. Stopping the control program still results in a large post-control outbreak and a divorce effect. **(b) Reactive Control for Seasonal SIR.** A fixed length (1/12 yr.) control is implemented to control an endemic disease (parameterized as in Figure 2(a)) once prevalence rises above a threshold (200 individuals in a population of 1 million). This stops the large early season outbreaks seen in the pulsed control, however the frequency of treatment increases as the susceptible population grows. Stopping the control program results in a large outbreak and divorce effect. **(c) Informed Control in seasonal SIR model.** The first control period occurs at time 0. The beginning of the next control period is decided at the end of the previous control period, and is the day (allowed to be up to a maximum of 365 days later) that will result in the smallest divorce effect if control was stopped after that period. This plan finds that it is optimal to perform the first few treatments relatively quickly, then to perform subsequent treatments during the peak in prevalence. We see that this is capable of nearly eliminating the Divorce Effect, but there is only a minimal benefit to the control, with large yearly outbreaks.

The reactive control has a similar effect following the initial control period, however it results in ever more rapid need for control, exhausting all 12 months of treatment in the first four years for both the directly transmitted infection (Figure 4(b)). We see that while this results in a lower RCI during the control program, it results in an even larger post-control outbreak and a larger maximum RCI for both transmission pathways.

Intuitively, Figure 2(c) suggests choosing a time period to implement the control that will minimize the divorce effect. To do this, we implement a third method which optimally chooses the time at which to begin the next control period. For this, we simulate the first one month control period, beginning at time 0. Then we run simulations with the next one month control beginning on all possible days over the next 365 days after the control ends, choosing the day that results in the lowest maximum RCI over the next decade, simulating through that control period, and repeating. This plan results in implementing the first three control periods in rapid succession and the remainder after the peak of an outbreak, when the control will have the least effect on transmission (Figure 4(c)), minimizing the magnitude of the divorce effect albeit at the cost of not providing significant protection against the infection. This result, along with other earlier results, suggests that the divorce effect is unavoidable and the potential for a divorce effect will continue to grow in magnitude unless the control is maintained for decades, regardless of the timing of the treatments. While it may not be possible to eliminate the divorce effect for relatively short controls, it may be possible to extend programs without worsening the divorce effect and to minimize the divorce effect by carefully choosing the timing of the end of the control program once cessation becomes necessary.

In the case of host-vector transmission, the yearly control successfully suppresses the infection for the first 1.5 years, however the population begins to experience outbreaks during what was traditionally the off-season. After the control program is ended, the population enters a period of larger outbreaks occurring every three years (Figure S15(a)). The reactive control sees a similar result as the directly transmitted disease, with all twelve treatments used in the first 4 years (Figure S15(b)). For the third method, the optimal plan was to wait the maximum amount of time to deploy the control (Figure S15(c)). This is likely due to the peak of on outbreak not occurring within a year of the end of treatment in the seasonal host-vector model.

## Additional Results

Results for additional models, along with an analytical approximation to the magnitude of the divorce effect are included in the supplemental information.

## Discussion

It has long been appreciated that non-immunizing control measures deployed against endemic infections will result in a large short-term reduction in prevalence but will lead to a reduction in herd immunity, leaving the population at risk of large outbreaks after the cessation of control. Here we have shown, in quite general settings, that these outbreaks can be so large as to increase, counting from the time that control started, the total incidence of infection above what would have occurred if no control had been used—a result we call the divorce effect. This represents a failure for control of the worst kind, namely a control that increases the total incidence of the infection. Unfortunately, many commonly used disease control plans rely on temporary non-immunizing controls, meaning that populations may be left at risk of the divorce effect once the control measure is ended.

Controls that do not confer immunity—including isolation, use of drugs as a prophylaxis or to shorten duration of infectiousness or behavioral changes such as social distancing—are often deployed in epidemic settings, particularly for new pathogens for which a vaccine is unavailable, but may also be used to blunt seasonal outbreaks of endemic diseases. In these endemic settings, we have shown that it is important to weigh any potential benefit from these controls against the risk of post-control outbreaks and the divorce effect. While there are timeframes over which a temporary non-immunizing control has benefits, the severity of the post-control outbreak that results in the divorce effect will risk overwhelming even well-maintained healthcare systems.

Vector-borne infections represent the most common situation in which non-immunizing controls are regularly used against endemic diseases, e.g. insecticide spraying to combat seasonal dengue outbreaks. The honeymoon effect predicts that insecticides can provide short-term benefits in endemic settings but that the additional benefit of continued spraying will decrease over time due to the accumulation of susceptibles (i.e. depletion of herd immunity) that results. Indeed, Hladish *et al.* [17] saw precisely these effects using a detailed agent-based model for dengue control that employs indoor residual spraying. Cessation of spraying will be expected to lead to large post-control outbreaks: again, Hladish *et al.*’s model exhibited annualized incidence of 400% compared to the uncontrolled baseline setting in certain years. Here, we examine the divorce effect directly and show that they are not specific to a host-vector model and that if the control is not maintained indefinitely, or at least for a few decades, the damage of the divorce effect can quickly outweigh the short-term benefits. Further, programs implementing insecticides may be intended to be indefinite, but the evolutionary pressure imposed can result in the rapid and unpredictable evolution of resistance. Without proper monitoring, this could result in an increase in total incidence due to the divorce effect before officials realize that resistance has developed. While insecticides, and other non-immunizing controls, will, and should, continue to play an important role in epidemic settings, where herd immunity is negligible, the results of this study raise important questions about their use in combating endemic infections.

In some instances, control measures are deliberately transient in nature, such as field trials for assessing the impact of proposed novel control methods, e.g. a review of field trials of dengue vector control showed they lasted between 5 months and 10 years [20]. Multiple year field trials such as these can result in considerable build-up of the susceptible population, meaning consideration needs to be given to the consequences of this accumulation and the potential for large outbreaks to occur in the wake of cessation of the trial. If our results are validated, they must be factored not only into the design of such trials but also into the informed consent process for trial participation, with participants made aware of the risk of the divorce effect and plans put in place to provide a reasonable level of protection during and following the study. As we have shown, these outbreaks can occur months or even many years later, and while disease incidence would be observed closely during the trial, our results argue that monitoring should continue for an appropriate length of time following the cessation of control. Furthermore, we emphasize that the epidemiological consequences of the honeymoon effect—specifically the relative ease of reducing incidence for an infection near endemic equilibrium—must be kept in mind when interpreting the results of such trials. Together, these dynamical effects argue that susceptibility of the population to infection should be monitored together with incidence to fully assess the impact and effectiveness of the control.

Additional concerns are raised when an endemic and an epidemic infection share the same transmission pathway (e.g. *Aedes aegypti* vectoring both dengue and Zika). Emergency control against the epidemic infection also impacts the endemic infection, leading to the potential for the divorce effect to occur in the latter if the control is ceased once the epidemic has subsided. It may be that policy makers have to choose to allow an epidemic of a highly publicized, but low risk, epidemic in order to maintain immunity levels of another lower profile, but more dangerous, disease. On the other hand, if the risk due to the epidemic is sufficiently high, it may still be advantageous to use the control, however the risks need to be carefully compared and an informed decision, that accounts for the divorce effect, needs to be made.

While transient non-immunizing controls are common and provide opportunities to observe the divorce effect, researchers tend to focus on prevalence or incidence over short periods of time and not cumulative measures such as CI or relative measures such as RCI or CA, which would expose the divorce effect. Even when relative measures are used, such as Hladish et al. [17], the time frame over which incidence is compared can have a drastic effect on the interpretation of the result. The divorce effect is an easily missed phenomenon, even when examining models that lack much of the real-world complexity, but real-world data comes with a myriad of other problems. Often the divorce effect may occur when the system is poorly monitored, as with field trials and unintentional control, in systems that, like dengue, have large year-to-year variation, or in systems where the failure is associated with other confounding socio-economic events such as war or natural disaster, resulting in data that is either scarce or difficult to interpret. The divorce effect may become more apparent in coming years, though, as mosquito control is lessened following the end of the Zika epidemic, allowing for a rebound in dengue in areas such as South America, and as insecticide resistance problems continue to grow.

Careful thought should be given to whether or not it is appropriate to begin new programs that rely on non-immunizing controls in endemic settings. This is an inherently complicated decision that must take into account numerous factors, both scientific and sociopolitical, but, in light of our results, policymakers should carefully weigh the risks of the divorce effect against other factors, e.g. imminent approval of a new vaccine or political pressure, before implementing disease management plans that rely on non-immunizing controls. Further, it is important that when non-immunizing controls are included in these management plans that they are not considered possible solutions but instead stop-gaps, and emphasis is placed on the development of vaccination as opposed to the indefinite continuation of the program.

Currently, control of endemic diseases worldwide, especially vector-borne diseases, relies heavily on non-immunizing controls such as insecticide. Policy makers should begin developing exit plans for these disease management programs —guidelines for safely ending the program when it becomes clear that indefinite maintenance is unlikely, which should be designed to minimize the impact of the divorce effect. In this paper, we have shown three possible designs for exit plans that could minimize the divorce effect. However, none of these designs were capable of eliminating the divorce effect. Our results suggest there is an inherent cost associated with the loss of immunity resulting from these programs.

## Acknowledgements

We thank Fred Gould, Sumit Dhole, Michael Vella, Christian Gunning, Jennifer Baltzegar, and Jaye Sudweeks for helpful discussion.

