## Supplemental Information for "After the honeymoon, the divorce: unexpected outcomes of disease control measures against endemic infections"

**Table of Contents:**


| | Page |
|---|---|
| **SIR Model Additional Figures** | **2** |
| **Seasonal SIR Model Additional Figures** | **8** |
| **Host-Vector Model Figures** | **11** |
| **Seasonal Host-Vector Model Additional Figures** | **13** |
| **Mitigating the Divorce Effect Additional Figures** | **15** |
| **SIR Model with Changing Population Size** | **18** |
| **SIR Model with Vaccination** | **19** |
| **SIRS Model** | **20** |
| **HIV Model** | **21** |
| **Age-structured Model with Realistic Mixing** | **23** |
| **Analytical Approximation** | **24** |
| **Sensitivity to Background Force of Infection** | **27** |

**SIR Additional Figures:**

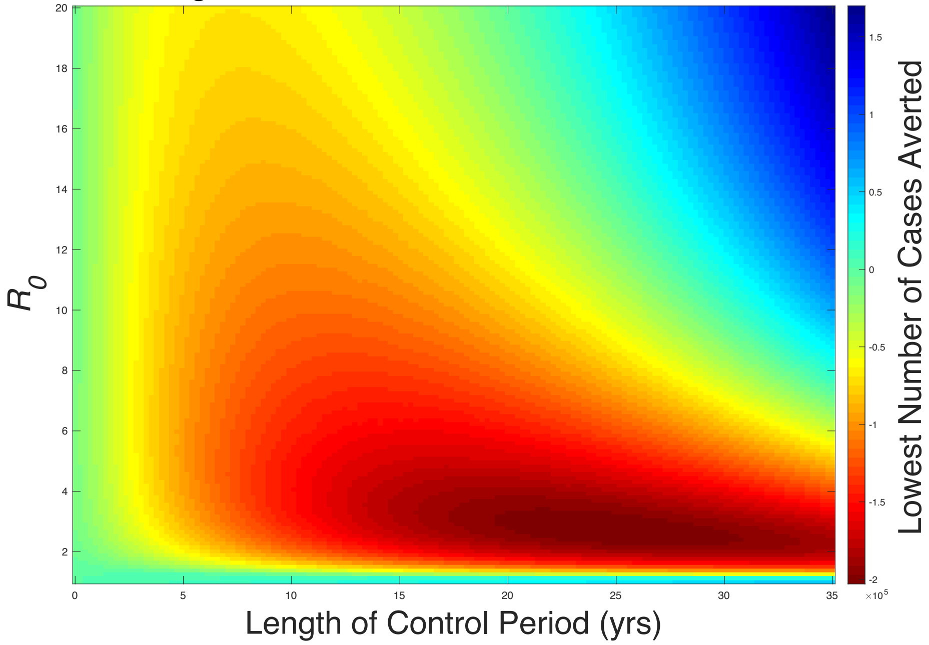


**Figure S1: Divorce Effect in Terms of Cases Averted.** All parameters the same as in Figure 1(b). When measuring the success of a control program in terms of Cases Averted as opposed to RCI, the overall results are retained, with negative values of cases averted corresponding to an RCI > 1. Parameters are as in Figure 1(b) for comparison.

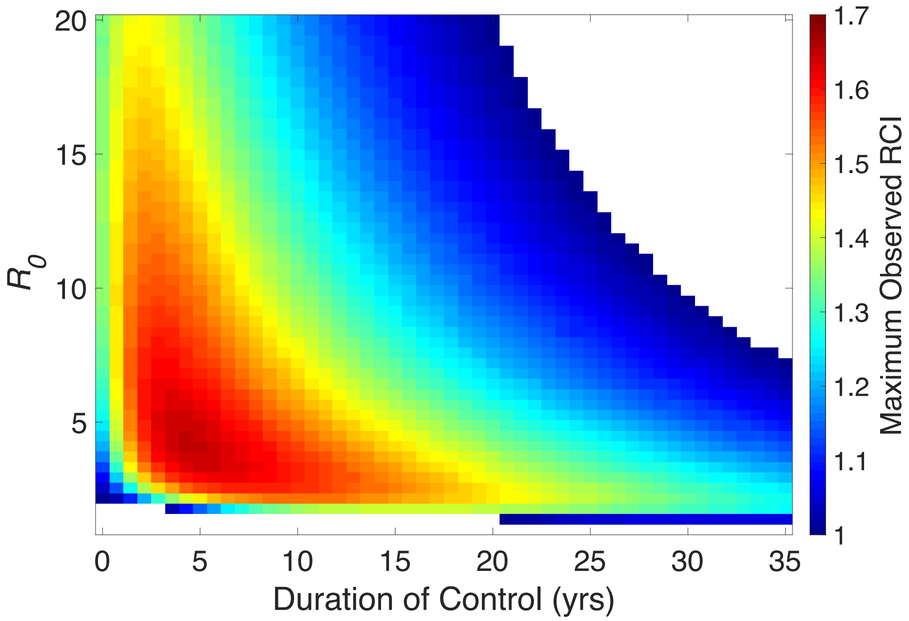


**Figure S2 Heat map of Divorce Effect from a Control that Increases Recovery Rate.** All parameters are as in Figure 1(b). The Divorce Effect is still observed if the control increases the recovery rate, $\gamma$, as opposed to decreasing the transmission parameter. All parameters the same as in Figure 1. Control increases $\gamma$ to 730 (previously 73 /year).

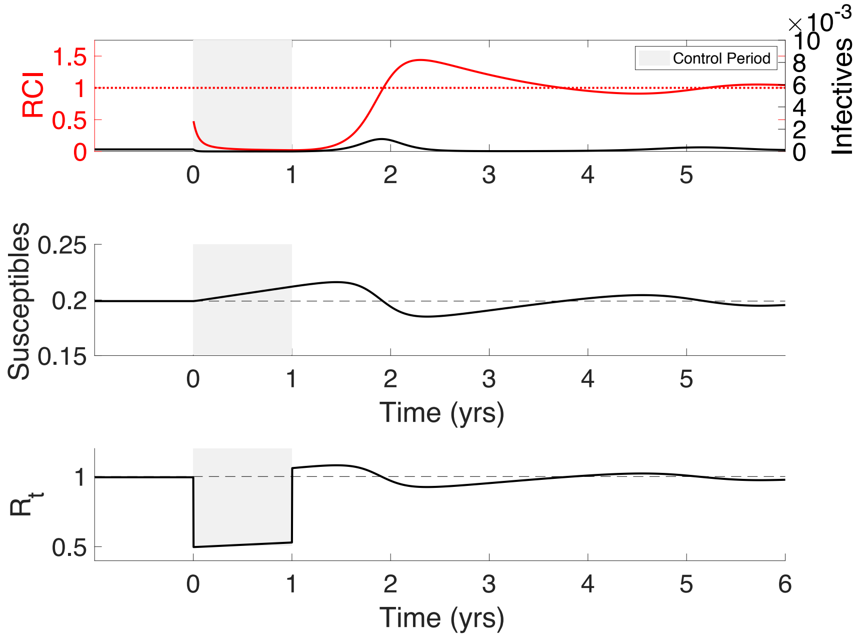


**Figure S3: Time-series showing the divorce effect in non-seasonal SIR model.** Figure corresponds to Figure 1(a) of the main text. Beginning at time zero, a year-long 50% reduction in the transmission parameter of an endemic infection ($R_0$=5) reduces prevalence of the infection to near zero for the length of the control, where it remains until time 1.5 yrs, at which point a large post-control outbreak occurs. RCI falls towards zero as prevalence remains low, but the post-control outbreak is large enough to bring RCI well above 1 (approx. 1.6). Panel 2 shows that the susceptible population begins to rise during the control period and continues until the outbreak depletes the susceptible population. Likewise, the reproductive number at time t, $R_t$, begins to rise during the control period. Once the control is released and the transmission rate retains its original value, $R_t$ increases above one and continues to grow until an outbreak occurs.

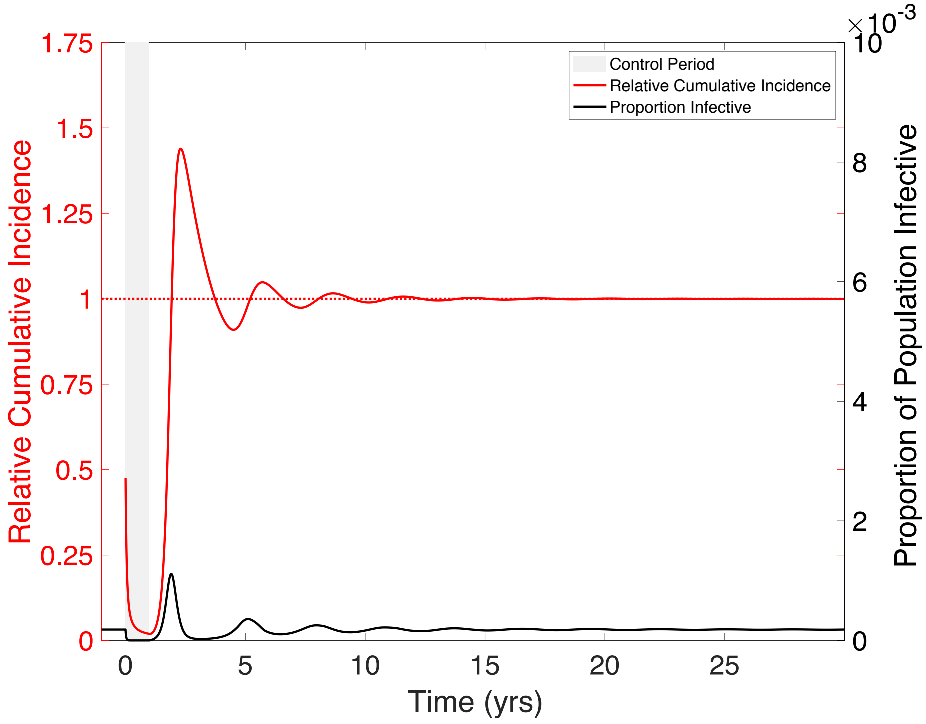


**Figure S4: Long-term time-series for the divorce effect in an SIR model.** Figure corresponds to Figure 1(a) of the main text. Following a one year control in which the transmission parameter is reduced by 50%, the host population continues to experience outbreaks that bring RCI above one until the infection approaches the endemic state and RCI approaches one.

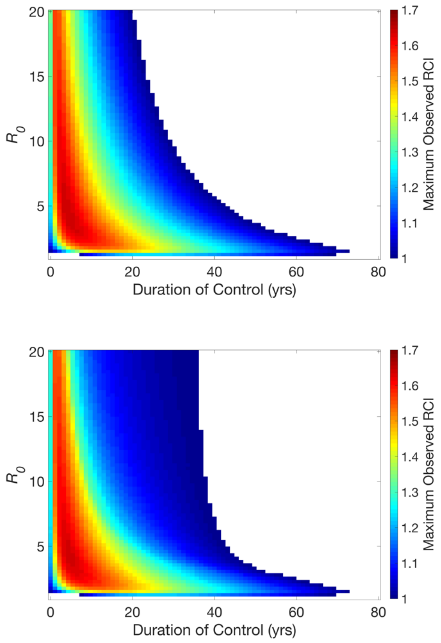


**Figure S5: Heat maps of Divorce Effect in SIR model for Different Strengths of Control.** We see that the Divorce Effect occurs in a significant area of the parameter space with controls that reduce the transmission parameter, $\beta$, by 100% (a), 75% (b), and 50% (c). In each case, the only way to avoid the divorce effect is to maintain control for more than 20 years (or approximately 40 years in the case of 50% control).

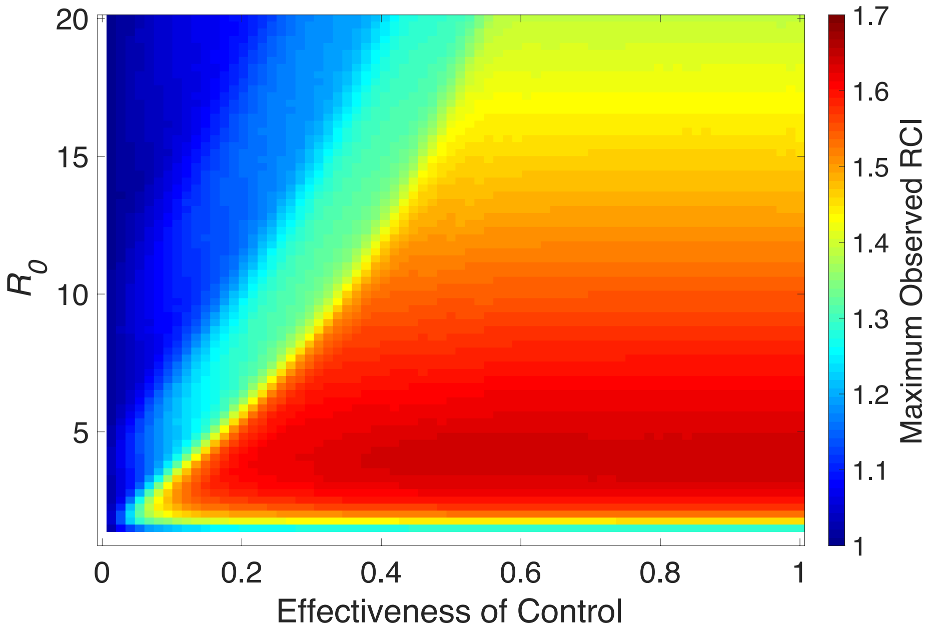


**Figure S6: Maximum RCI for a given effectiveness of control and $R_0$.** All controls are assumed to last for 1 year. The maximum RCI is found as the maximum observed RCI within 25 yrs after the end of a control that is between 0% and 100% effective for an infection with an $R_0$ of between 0 and 20. The areas of lowered maximum RCI result from outbreaks due to honeymoon effect outbreaks deplenishing the population of susceptible individuals before the control periods end.

### Seasonal SIR Model Additional Figures

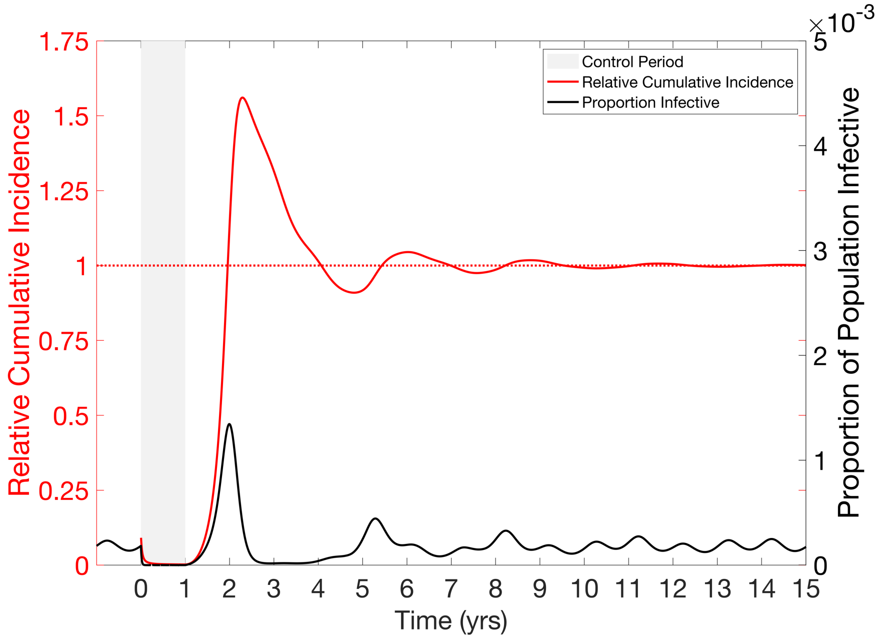


**Figure S7: Long-term time-series for the divorce effect in a seasonal SIR model.** Following a one year control in which the transmission parameter is reduced by 50%, the host population continues to experience outbreaks that bring RCI above one until the infection approaches the endemic state and RCI approaches one.

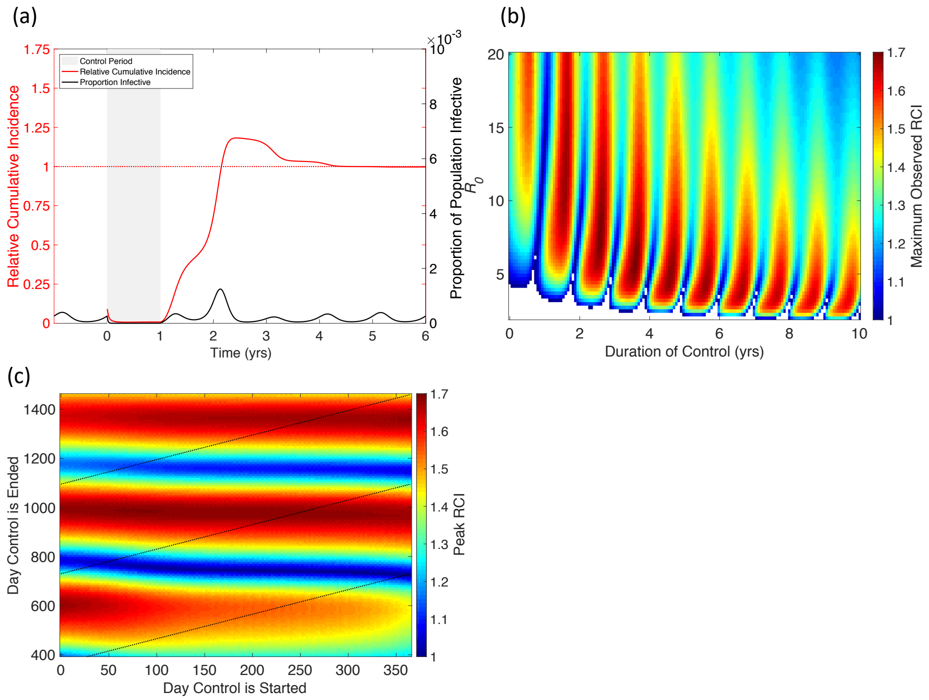


**Figure S8: The divorce effect in the seasonal SIR model with higher seasonality.** $\beta_1 = .1$, $I_b = 10$, all other parameters as in Figure 2. Here a higher value of $I_b$ is taken to adjust for the higher seasonality of the model. **(a) Typical time-series showing the divorce effect.** Beginning at time zero, when the transmission parameter is at its maximum, a year-long 90% reduction in the transmission rate of an endemic infection ($R_0$=5) is implemented at the beginning of a seasonal outbreak and reduces prevalence of the infection to near zero for the length of the control. Following the end of the control, a small late season outbreak occurs, followed by a large outbreak during the next season. RCI falls towards zero as prevalence remains low while the control is in effect, rises slightly during the small late season outbreak, and rises above 1 during the large outbreak the following year. **(b) Magnitude of divorce effect in terms of relative cumulative incidence (RCI).** Maximum RCI is found as the highest value RCI observed within 25 yrs following a 100% effective control of an infection with 1<$R_0$<20 and lasting between 1 month and 20 years. RCI>1 indicates the divorce effect and we see that divorce effect occurs in most of the parameter space. Unlike the SIR model (Figure 1), the magnitude of the divorce effect is not solely dependent on $R_0$. **(c) Effect of timing on the magnitude of the divorce effect.** Maximum RCI is the highest RCI observed within 25 yrs following a 100% effective control of an infection with $R_0$=10 beginning and ending on specified days. Dashed lines represent controls lasting either 1, 2, or 3 years. Maximum RCI is most sensitive to the day the control is ended, moderately sensitive to the day it is started, and only slightly sensitive to the length of the control. This is due to the timing of the end of the control determining the timing of the outbreak. We also see that continuing the control for another year has little impact on the magnitude of the divorce effect.

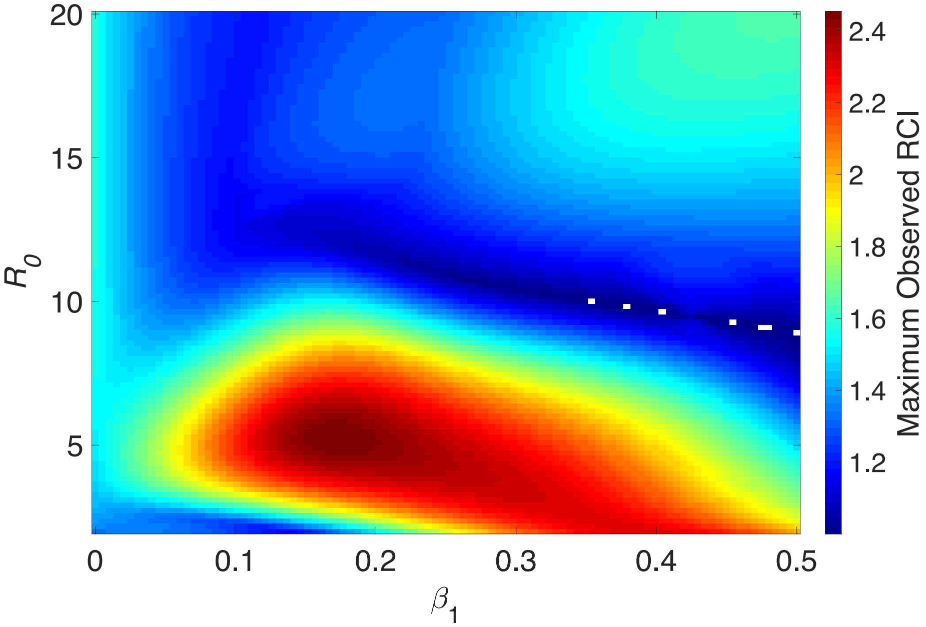


**Figure S9: Maximum RCI for a given basic reproductive number and seasonality in the directly transmitted model.** All controls are assumed to increase the vector mortality rate by 100% and to last for 1 year. The vector reproductive rate is assumed to have some average rate, *r*, and some level of seasonality ($r_s$). The maximum RCI is found as the maximum observed RCI within 25 yrs after the end of a control. Here we see that the divorce effect is present throughout most of the parameter space.

Non-Seasonal Host-Vector Figures

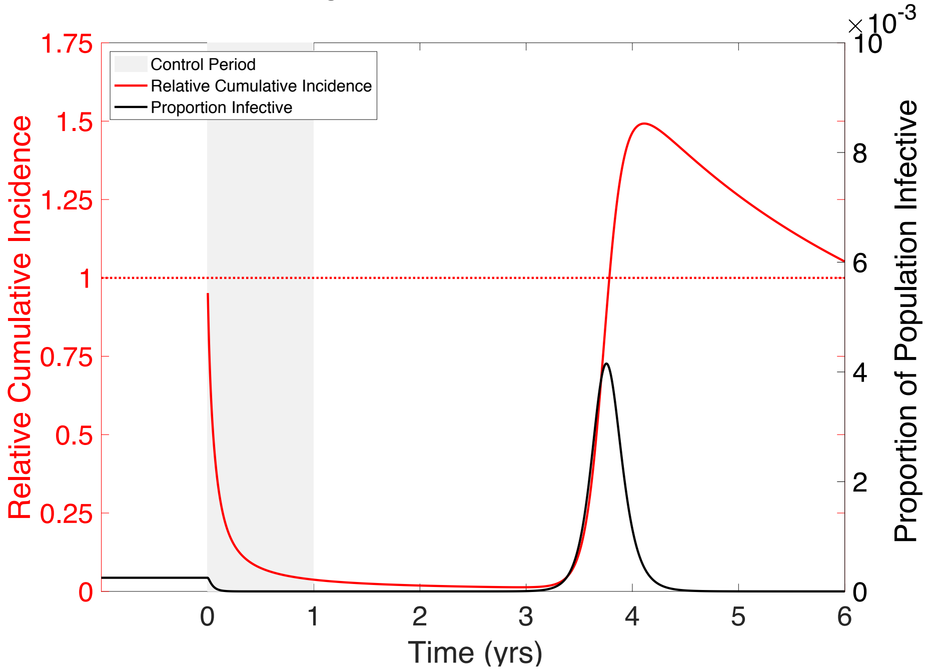


**Figure S10: Time-series of the divorce effect in a non-seasonal vector-borne infection.** Figure shows the result of the non-seasonal host-vector model given in main text. Following a year of control, against an endemic infection ($R_0 = 5$), in which the vector lifespan is reduced by 50% ($\delta = 73/\text{year}$ increased from $\delta = 36.5/\text{year}$) incidence is reduced to near zero. After the control is stopped, we see a post-control outbreak in year 3, resulting in the divorce effect (peak RCI≈1.5).

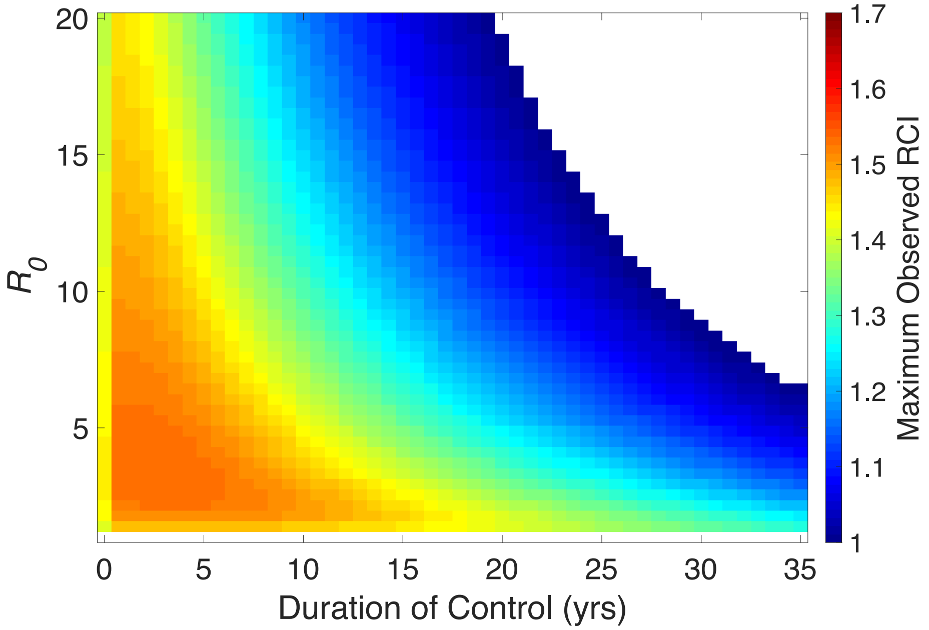


**Figure S11: Heat Map for non-seasonal Host-Vector Model** – Maximum RCI is the highest RCI observed within 25 yrs following control of an infection with $R_0$ = 5 beginning and ending on specified days. We see for the Host-Vector model that, much like the SIR model, the only way to avoid the divorce effect is to maintain control for more than 20 years. Parameters as given in the main text. Control decreases vector life-span by 50%.

### Seasonal Host-Vector Model Additional Figures

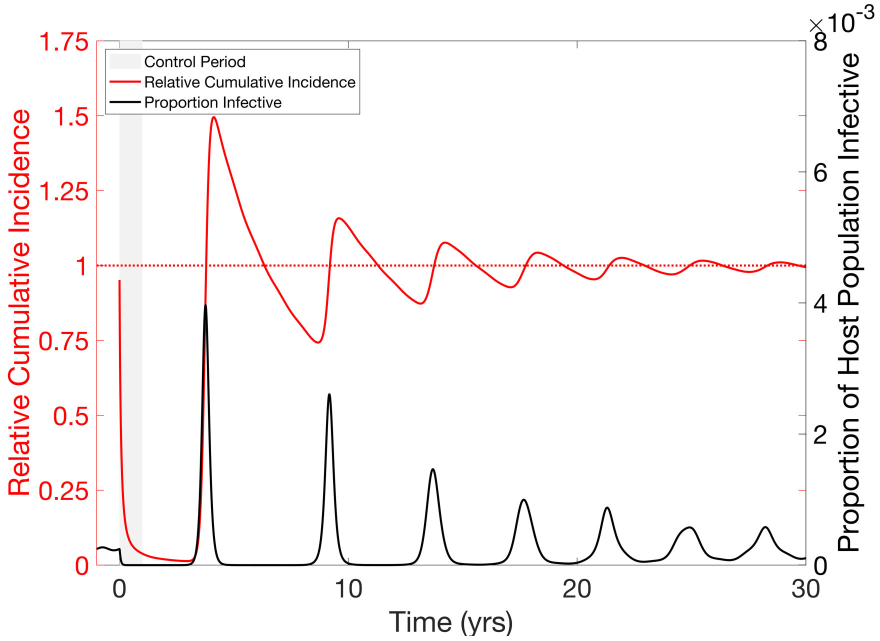


**Figure S12: Long-term time series for the divorce effect in a seasonal host-vector model.** All parameters are as in Figure 3 of the main text. Following a one-year control, a large outbreak occurs in year 3, that brings RCI above 1. This outbreak depletes the susceptible population, resulting in no outbreaks for the next five years. Each subsequent outbreak is sufficiently large to bring RCI above 1. Around year 30, the system is still experiencing larger than normal outbreaks, bringing RCI slightly above 1.

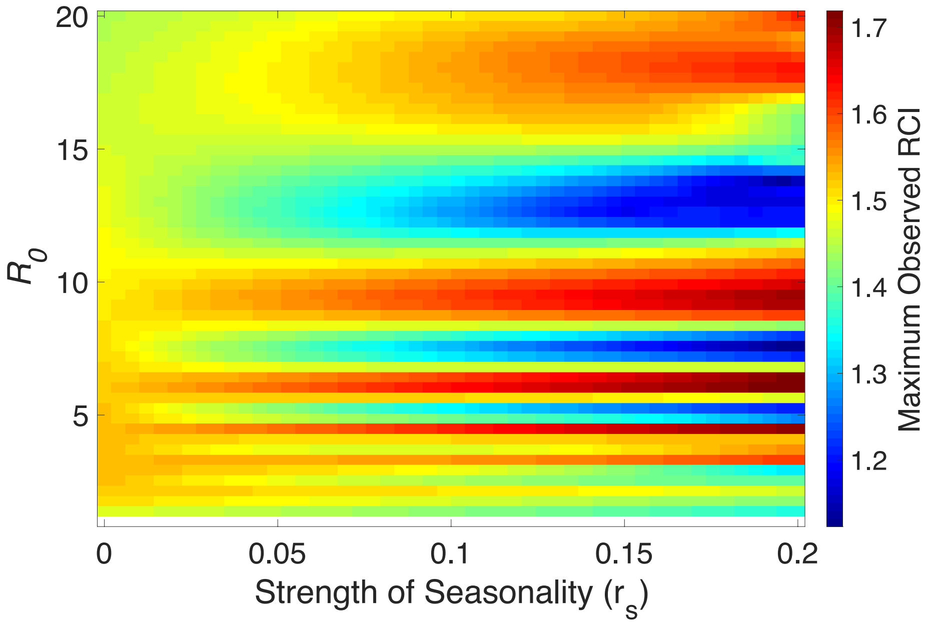


**Figure S13: Maximum RCI for a given basic reproductive number and seasonality in the host-vector model.** All controls are assumed to increase the vector mortality rate by 100% and to last for 1 year. The vector reproductive rate is assumed to have some average rate, *r*, and some level of seasonality ($r_s$). The maximum RCI is found as the maximum observed RCI within 25 yrs after the end of a control. Here we see that the divorce effect is present throughout the parameter space.

**Mitigating the Divorce Effect Additional Figures**

(a)

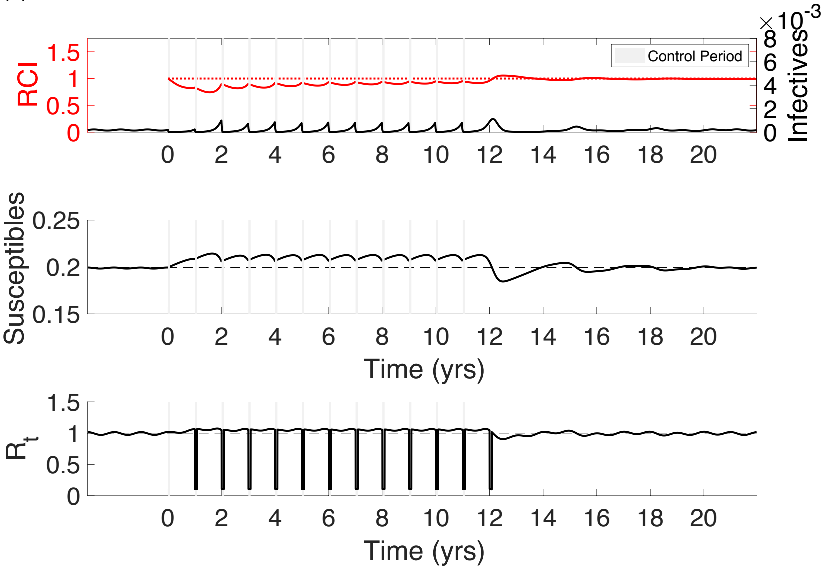


(b)

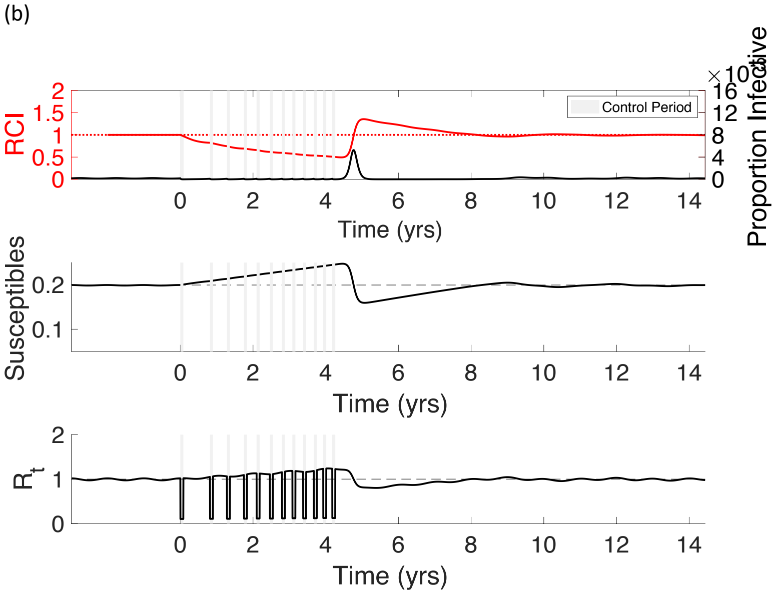

(c)

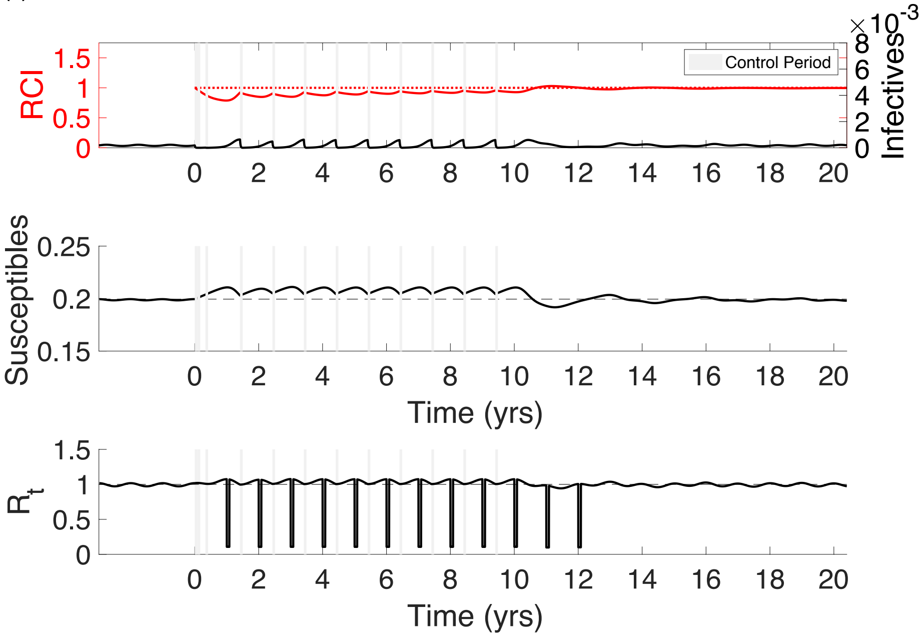


**Figure S14: Suggested techniques for mitigating the Divorce Effect in the seasonal SIR model**. In the pulsed (a), reactive (b), and informed (c) techniques, we see the susceptible population begin growing with the first treatment and continue growing until the outbreak occurs after the 12th treatment, at which the susceptible population is quickly depleted. Likewise, the reproductive number, $R_t$, increases overall during this time with seasonal fluctuations, and reductions due to control periods.

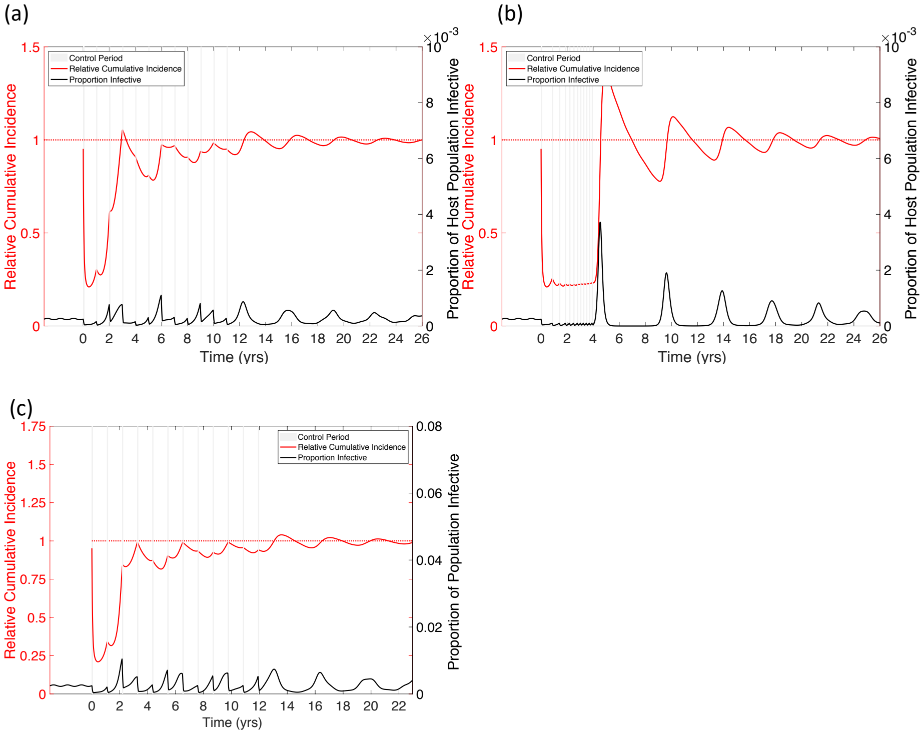


**Figure S15: Suggested techniques for mitigating the divorce effect in a seasonal host-vector model. (a) Pulsed control for Seasonal SIR model.** Control ($\sigma = 1$) occurs yearly at a fixed time (when $R_0$ is highest) for a fixed time (1 mo.). The control is effective at stopping the outbreak the first year, but seasonal outbreaks in subsequent years are of varied magnitudes due to the susceptible population being depleted in some years and replenished in others. An outbreak in year 2 is large enough to result in the divorce effect. Stopping the control program results in a large post-control outbreak and a divorce effect. **(b) Reactive Control for Seasonal SIR.** A fixed length (1 mo.) control is implemented once prevalence rises above a threshold (200 individuals in a population of 1 million). This stops the large outbreaks seen in the pulsed control, however the frequency of treatment increases as the susceptible population grows, and all treatments are depleted within the first four years. Stopping the control program results in a large outbreak and divorce effect. For all panels twelve 1 mo. controls are used to be consistent with the 1 yr. controls used in other figures. **(c) Informed Control in seasonal Host-Vector model.** Control works by increasing vector mortality by 100% for 1 month. The first control period occurs at time 0. The beginning of the next control period is decided at the end of the control period, and is the day that will result in the smallest Divorce Effect if control is stopped after that period (a maximum of 1 year between treatments). We see that this is capable of nearly eliminating the Divorce Effect, but there is only a negligible benefit to the control, with large yearly outbreaks. Importantly, this plan recommends waiting the full year, suggesting that the optimal timing of the next treatment may occur after this period.

### Additional Models:

#### SIR Model with Changing Population Size

We assume a population similar to the SIR model in the main text, with the exception that the per capita birth and death rate are allowed to differ. We have

$$\begin{aligned}\dot{S} &= bN - \mu S - \beta\frac{S(I + I_b)}{N}\\ \dot{I} &= \beta\frac{S(I + I_b)}{N} - (\gamma + \mu)I\\ \dot{N} &= (b - \mu)N.\end{aligned} \qquad \text{(S1)}$$

Here $b$ is the per capita birth rate and μ is the per capita death rate. For illustration, we take two values of $b$, $b = 1.25\mu$ and $b = .75\mu$, corresponding to 25% population growth or reduction per year. While this is an extreme case, we expect that any effect on the magnitude of the divorce effect would most likely be seen in the extremes. Since the endemic equilibrium is not well defined for a changing population, we simulate the population for a thousand years before starting control. Initial values of $N$ were chosen so that at the end of the thousand years, the population size was $1 \times 10^6$. We see that the growth (Figure S16a), or decline (Figure S16b), of the population does not eliminate the divorce effect, but does affect the magnitude and timing of the post-control outbreak, with a larger and earlier post-control outbreak in the growing population due to a larger number of susceptible individuals being born.

(a) (b)

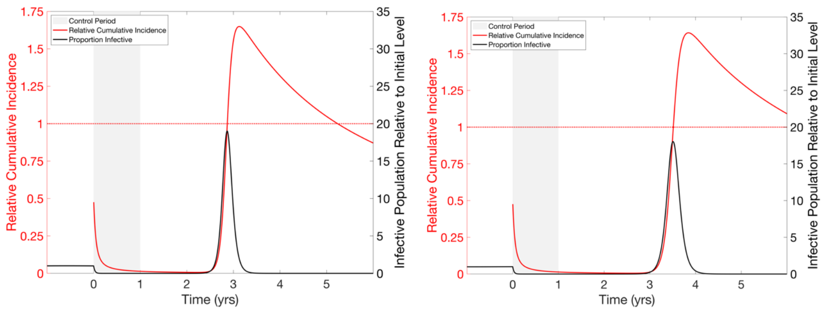


**Figure S16: The divorce effect in the SIR model with a (a) growing and (b) shrinking population.** The population is (a) growing or (b) shrinking during the simulation at a rate of 25% per year. Beginning at time zero, a year-long 50% reduction in the transmission parameter of an endemic infection ($R_0$=5) reduces prevalence of the infection to near zero for the length of the control, where it remains until time 2.5 yrs (a) or 3 yrs (b), at which point a large post-control outbreak occurs.  RCI falls towards zero as prevalence remains low, but the post-control outbreak is large enough to bring RCI well above 1 (peak RCI = 1.67 and 1.64 for the growing and shrinking population, respectively). Note that, because of the changing population sizes within and between graphs, prevalence of infection is plotted on a relative scale on both graphs.

**SIR Model with Vaccination**

To model vaccination against infection, we assume that some portion, $v$, of births enter the recovered class instead of the susceptible class, while all other dynamics proceed similarly to the SIR model (Equations S2).  For illustration, we take $v = .5$ and assume the vaccination campaign lasts one year before being discontinued.  We see that during the control period the proportion of the population that is infective falls significantly more slowly than with transmission reduction (Figure S17).  Following the end of control, we see a series of post-control outbreaks that bring the infective proportion of the population above endemic levels, but they are not large enough to bring RCI above 1.  This lack of divorce effect is due directly to the maintenance of population level immunity due to the vaccination, which keeps the susceptible population from being able to build sufficiently.  It is important to note that, as shown in Okamoto et al. [1], it is possible to see the divorce effect in combined controls that involve both immunizing and non-immunizing controls.

$$\begin{aligned} \dot{S} &= b(1 - v)(N - S) - \beta \frac{S(I + I_b)}{N} \\ \dot{I} &= \beta \frac{S(I + I_b)}{N} - (\gamma + \mu) I \end{aligned} \qquad \text{(S2)}$$

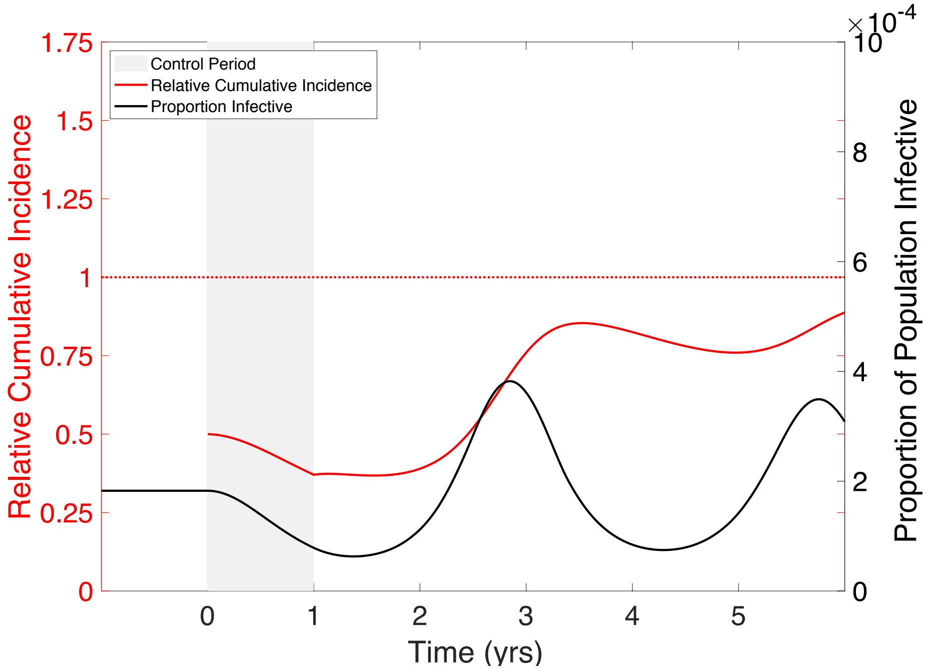

**Figure S17: Cessation of a vaccination program.** A vaccination program is put in place in which 50% of newborns are vaccinated for one year. All other parameters are as in Figure 1. The vaccination program is discontinued after the first year. While we see post-control outbreaks that bring incidence above the endemic level, they are not large enough to bring RCI above one, and RCI approaches 1 in the long run.

**SIRS Model**

We assume a well-mixed population with parameters defined as in the main text. However, instead of permanent immunity, we assume that immunity is lost at per-capita rate $l$, such that the average length of immunity following an infection is $1/l$ (Equation S3). For the sake of illustration, $l = 1/10 \text{ year}^{-1}$, corresponding to an average of 10 years of immunity following recovery.

$$\begin{aligned} S &= \mu(N - S) - \beta\frac{S(I + I_b)}{N} + lR \\ I &= \beta\frac{S(I + I_b)}{N} - (\gamma + \mu)I \\ R &= \gamma I - (\mu + l)R \end{aligned} \qquad \text{(S3)}$$

Similar to the SIR model, we see suppression of the infection for a period of time during and immediately following the control (Figure S18). A large post-control outbreak is seen about 3 months after the end of treatment. This outbreak is sufficiently large to bring the RCI above 1,

**Deleted:** ¶ ... [1]

to about 1.45, before the outbreak subsides and prevalence and RCI fall again. As the immune period following infection shrinks towards zero, the SIRS model approaches the behavior of an SIS model. This results in the magnitude of the divorce effect being reduced as the immune period, and the population of immune individuals, becomes smaller.

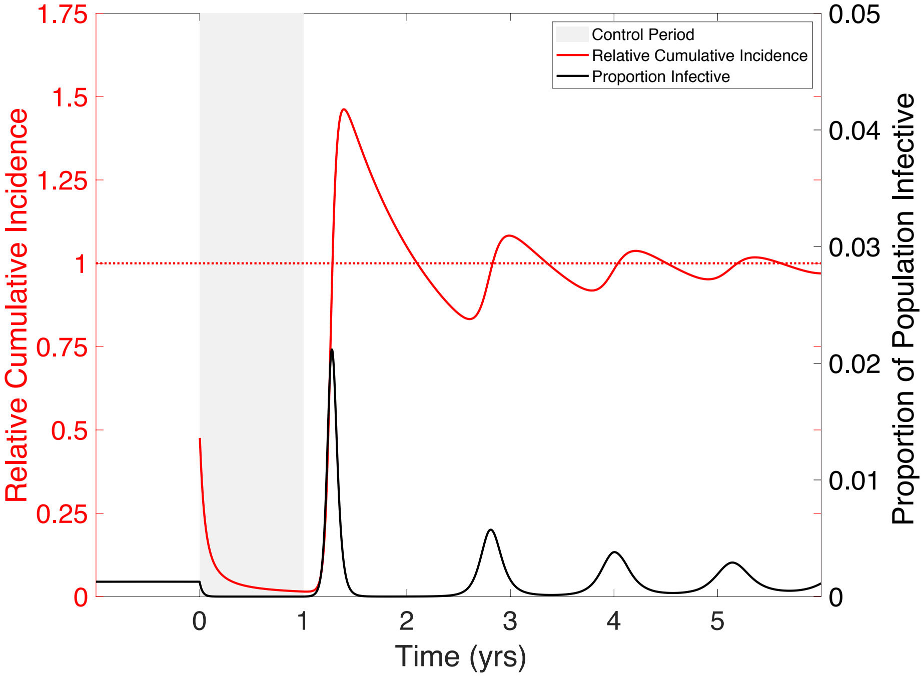


**Figure S18: Divorce Effect in an SIRS model.** Beginning at time zero, a year-long 50% reduction in the transmission parameter of an endemic infection ($R_0$=5) reduces prevalence of the infection to near zero for the length of the control, where it remains until time 1.5 yrs, at which point a large post-control outbreak occurs. RCI falls towards zero as prevalence remains low, but the post-control outbreak is large enough to bring RCI well above 1 (peak RCI is approx. 1.45).

**Within-Host Virus Dynamics (HIV) Model**

We examine the divorce effect in the model for the within-host dynamics of HIV presented in Rong and Perelson [2], with all equations and parameters taken directly from their text (Equations S4). Here, $T$ stands for the concentration of target cells, $L$ for latently infected cells, $T^*$ for actively infected cells, $V_I$ for infectious virions, and $V_{NI}$ for non-infectious (defective) virions. Parameter names and values are given in Table 1. Here, cumulative incidence is in terms of actively infectious T cells. We see that the divorce effect does occur following a 25 day treatment that has both a protease inhibitor and reverse transcriptase inhibitor with efficacies of 50% (Figure S19).

$$\begin{gathered} T = \lambda - d_T T - (1 - \epsilon_{RT}) k V_I T \\ L = \alpha_L (1 - \epsilon_{RT}) k V_I T - d_L L - aL \\ T^* = (1 - \alpha_L)(1 - \epsilon_{RT}) - \delta T^* + \alpha L \\ V_I = (1 - \epsilon_{PI}) N \delta T^* - c V_I \\ \dot{V_{NI}} = \epsilon_{PI} N \delta T^* - c V_{NI} \end{gathered} \tag{S4}$$

Deleted: ¶
¶ ... [2]

| Symbol | Parameter | Value |
|---|---|---|
| $\boldsymbol{\lambda}$ | Susceptible Cell Recruitment Rate | $10^{-4}$ mL$^{-1}$day$^{-1}$ |
| $\boldsymbol{d_T}$ | Susceptible Cell Mortality Rate | $.01$ day$^{-1}$ |
| $\boldsymbol{\alpha_L}$ | Fraction of Infections Resulting in Latency | .001 |
| $\boldsymbol{k}$ | Infection Rate Constant | $2.4\text{x}10^{-8}$mL day$^{-1}$ |
| $\boldsymbol{d_L}$ | Death Rate of Latent Cells | $.004$ day$^{-1}$ |
| $\boldsymbol{\alpha}$ | Latent Cell Activation Rate | $.1$ day$^{-1}$ |
| $\boldsymbol{\delta}$ | Death Rate of Actively Infected Cells | 1 day$^{-1}$ |
| $\boldsymbol{c}$ | Free Virus Clearance Rate | 23 day$^{-1}$ |
| $\boldsymbol{N}$ | Burst Size | 3000 |
| $\boldsymbol{\epsilon_{RT}}$ | Efficacy of Reverse Transcriptase inhibitors | .5 |
| $\boldsymbol{\epsilon_{PI}}$ | Efficacy of Protease Inhibitors | .5 |

**Table 1:** Parameter values for the HIV model.

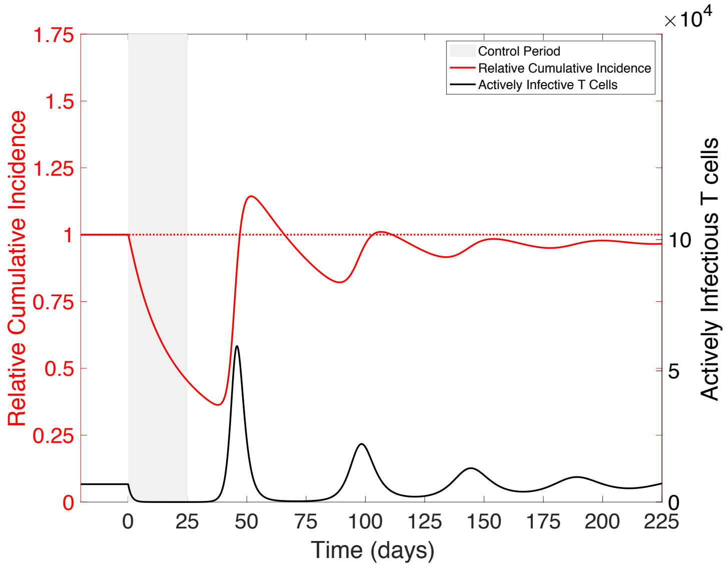


**Figure S19: Divorce effect in the within-host virus dynamics (HIV) model.** Beginning at time zero, a 25 day treatment occurs using a drug that combines a protease inhibitor and a reverse transcriptase inhibitor, both with 50% efficacy. This successfully reduces the infectious T cell count to near zero during and immediately following the treatment period. After the end of treatment, we see a transient increase in infectious T cells, bringing the relative cumulative incidence of T cell infection above one (max RCI≈$1.14$). After years, RCI eventually approaches 1 from below.

**Age-structured Model with Realistic Mixing**

Here we show the presence of the divorce effect in an age-structured model with realistic mixing between groups. This model, and code, is from a tutorial given by Aaron King and Helen Wearing [3]. We assume that there 30 age-groups, with ages 0-19 occurring as single year age groups, 20-75 as 5 year age groups. Transitions between compartments occur according to Equation S5, note that we use ∘ to denote elementwise multiplication. In which *A* is a matrix describing transitions between age classes, e.g. aging and deaths, *b* is a matrix describing births with a constant birth rate as its first element and zeros everywhere else. New-born susceptibles enter the youngest age class at a rate of *b* = 100/year, movement between the age classes takes on average 1 year for ages 0-20, 5 years for ages 21-75, and death occurs at a constant rate in the last age class, occurring on average after 15 years. $S, I,$ and $R$ are vectors containing the numbers of individuals of each age class that are susceptible, infective, or immune, respectively. $\beta$ is a matrix containing the transmission parameters for infection occurring within and between age classes, and is constructed by taking a matrix of age-specific contact rates and multiplying it by a constant rate of infection per contact. This contact network is based on [4] and freely available online, and the constant rate of infection per contact chosen so that $R_0 = 5$. $\gamma$ is a vector containing the rate of recovery of individuals in each age class, but is assumed to be constant across all age classes and is the same as the main text $(\gamma = 73\ /\text{year})$. Control works, as in the SIR model, by reducing the transmission parameter by 50% and lasts one year.

$$\begin{aligned} \dot{S} &= -\beta I \circ S + AS + \mathrm{b} \\ \dot{I} &= \beta I \circ S + AI - \gamma I \\ \dot{R} &= AR + \gamma I \end{aligned} \qquad \text{(S5)}$$

We see that, similar to the non-structured SIR model, there is a period of time, lasting about 4 years, in which RCI is falling, before a large outbreak brings RCI above 1 (Figure S20). Importantly, while the magnitude of the effect varies across groups, due to mixing, its presence does not.

**Deleted:** ¶ ... [3]

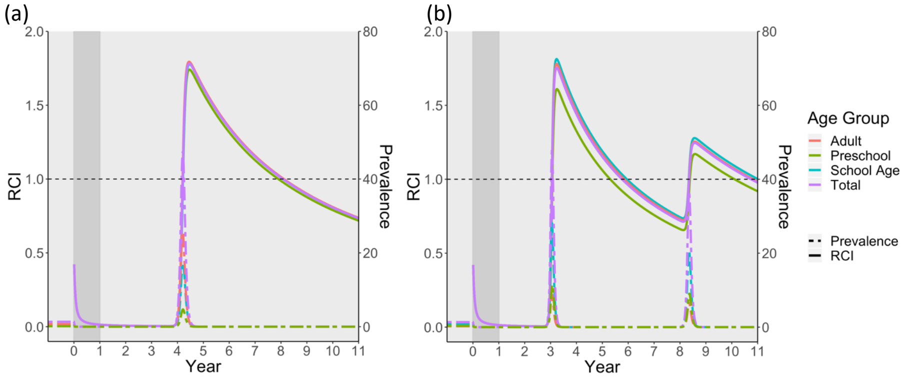


**Figure S20: Divorce effect in an age-structured SIR model with realistic mixing.** A control for a directly transmitted infection with $R_0$ = 5 (a) and $R_0$ = 15 (b) in a population of 9000 individuals is implemented at time 0, during which transmission is reduced by 50% for 1 year. At the end of the year, control is instantaneously removed. RCI quickly falls to 0 during the control period and remains there until a large outbreak in year 4 brings RCI up above 1 for all age groups. For the figure, prevalence, the number of individuals currently infective, (dashed lines) and RCI (solid lines) are shown for the total population, and age groups are shown aggregated into three groups: preschool (ages 0-5), school age (ages 6-18), and adult (ages 19+).

**Analytical Approximation:**
Here we describe a crude analytical approximation for the magnitude of the divorce effect in the simplest setting of a non-seasonal directly transmitted infection (i.e. the SIR model), and based on the well-known analysis of the size of an outbreak in a closed population [5,6]. We assume that the post-control outbreak occurs immediately following the end of the control period and that the outbreak happens instantaneously. Further, we assume that control is perfect, so that there are no new cases of infection during the control period, and that all individuals that are infective before the control begins recover by the end of the control period. When control begins, the population can be subdivided into individuals that are susceptible and those that have previously been exposed and will be immune when the control is ended. Assuming $R_0$>1, the numbers in these two groups are determined by the endemic equilibrium, where $S^* = N/R_0$ and $R^* = N(1 - 1/R_0)$. The number in the latter group decays exponentially due to mortality and the number of susceptibles grows at the same rate because of births (noting that the population size is taken to be constant). This gives the number of susceptible individuals at the time control ends, $t_{\mathrm{end}}$, as

$$S = N\left(\frac{1}{R_0} + \left(1 - \frac{1}{R_0}\right)\left(1 - e^{-\mu t_{end}}\right)\right) \qquad \text{(S6)}$$

Once the control is ended, the infection is assumed to be reintroduced immediately by a small number of infectious individuals and occurs instantaneously, meaning that demography does not affect the final outbreak size. This means that the post-control outbreak size, *Z*, can be found by solving the familiar transcendental equation:

$$Z = S\left(1 - e^{-R_0\left(\frac{Z}{N}\right)}\right). \quad \text{(S7)}$$

The post-control outbreak size is then compared to the cumulative number of infections that would be expected in the endemic case to find the predicted RCI (Equation S8).

$$\mathrm{RCI} = \frac{Z}{\mu N\left(1 - \frac{1}{R_0}\right)t} \quad \text{(S8)}$$

**Results of Analytical Approximation**

When compared to the simulations, our analytical approximation overestimates the magnitude of the divorce effect (Figure S21(a)). This is in direct contrast to simulations where the outbreak requires a long accumulation of infectives, often happens years later, and takes some time to occur. This approximation performs best in the most biologically relevant portion of parameter space ($R_0$<10 and control lasting less than 20 years), where the error is generally below 20% (Figure S21(b)), however it performs very poorly for extremely short durations of control.

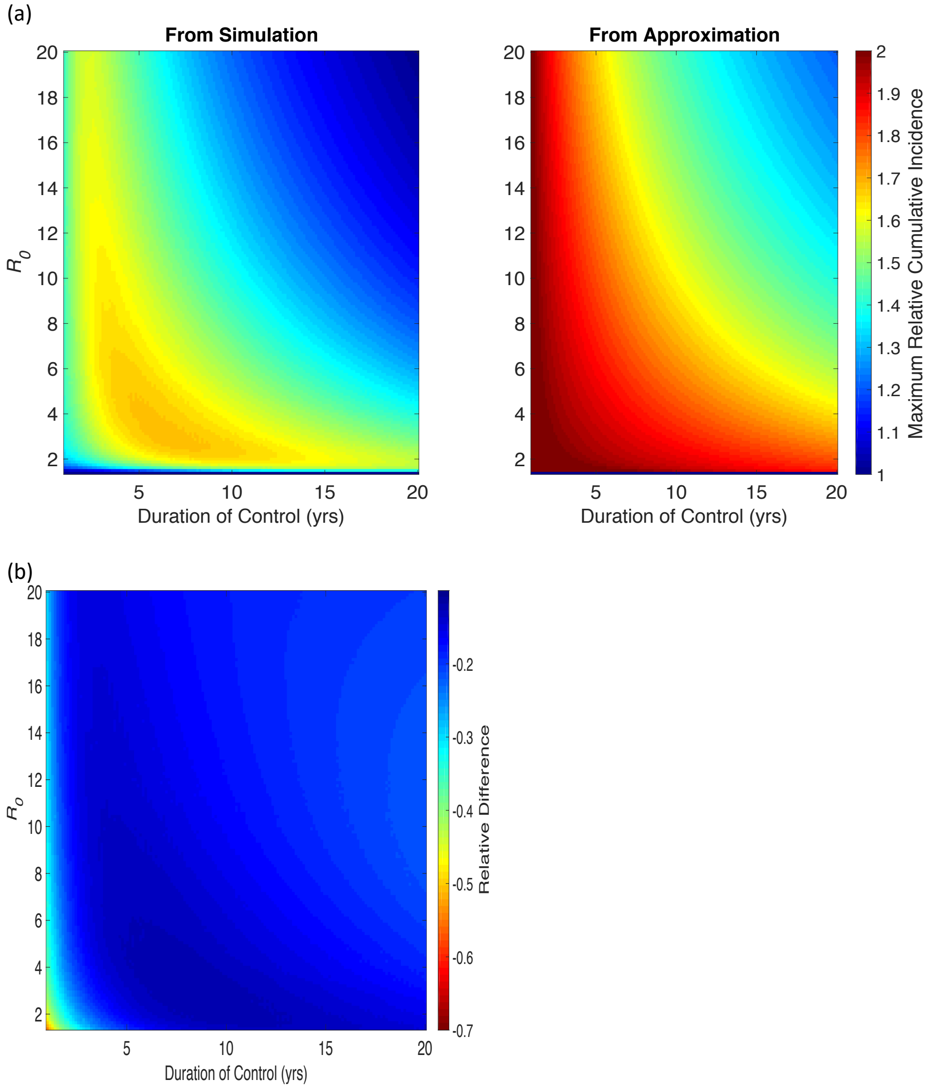


**Figure S21: Analytical approximation of the magnitude of the divorce effect. (a) Heat map of approximated and simulated magnitude of the Divorce Effect in terms of RCI.** The analytical approximation predicts the divorce effect for all controls lasting less than 20 years, similar to what is observed in the SIR model (Figure 1(b)). However, it over estimates the observed maximum RCI throughout the parameter space, and does so drastically for a short control in a system with $R_0$<3. **(b) Relative difference between the observed and predicted maximum RCI.** Calculated as (observed-predicted)/observed. The relative difference is small (<.25) throughout most of the parameter space except for short controls in systems with $R_0$<3.

### Sensitivity to Background Force of Infection

Deterministic compartmental epidemiological models suffer from the well-known weakness that the numbers of infectives can fall to arbitrarily low levels. To combat this, a background force of infection is often included in such models, representing infections due to contact with populations outside the focal population [7]. In our model, this process is accounted for by adding $I_b$ to the number of infectives in the transmission term. The background force of infection, which is taken to be small compared to the within-patch force of infection at the endemic state, ensures that there is a low level of transmission in the population, even as the number of infectives falls during the control period, and acts to reseed infection following control. In doing this, the background force of infection controls how quickly an outbreak will occur following the end of control, and hence can play an important role in determining the magnitude of the divorce effect. In general, a lower background force of infection means a later post-control outbreak, and often a larger divorce effect, while a higher background force of infection means an earlier post-control outbreak, less time for the build-up of the susceptible population, and a smaller divorce effect. These effects are most noticeable for a short-lived control. At a sufficient level, the background force of infection is large enough to drive the overall dynamics of the system, eliminating the divorce effect. When this occurs, the dynamics become driven by exogenous factors, similar to a sylvatic infection, reducing the importance of local infections. In addition to affecting the magnitude of the divorce effect, increasing $I_b$ increases the rate at which the system approaches its endemic equilibrium following the end of control. This results in subsequent outbreaks being increasingly diminished. For our manuscript, we choose to use a realistic value of $I_b = 1$ for our models, compared to an endemic level of 183 infective individuals for these parameter values in the nonseasonal model. Figure S22 shows that for values of $I_b$ that are sufficiently large to eliminate the divorce effect would require $I_b$ to be roughly the same size as the endemic infection level.

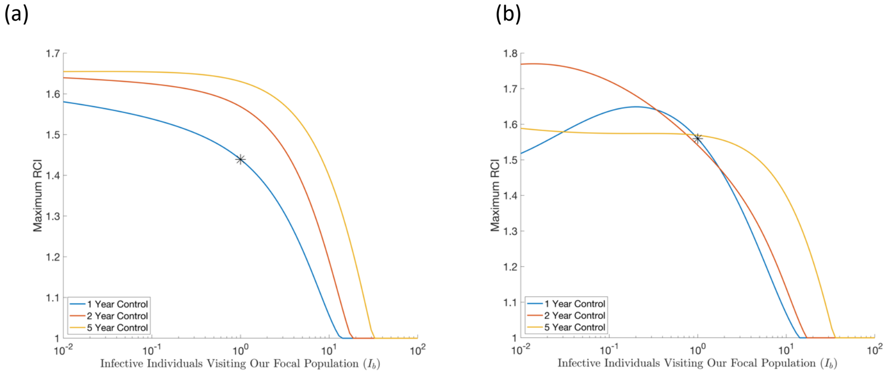


**Figure S22: Sensitivity of the magnitude of the divorce effect to the background force of Infection in the (a) non-seasonal and (b) seasonal SIR models.** Figure (a) is parameterized as in Figure 1(a) ($R_0 = 5$) and (b) is parameterized as in Figure 2(a) ($R_0 = 5$, $\beta_1 = .02$). All controls are assumed to last one, two, or five years, beginning at *t* = 0. We see that for a sufficiently high number of infective individuals visiting our focal population ($I_b$), the divorce effect is eliminated. We choose a seemingly realistic value of $I_b = 1$ (stars) for our models, compared to an endemic level of 183 infective individuals for these parameter values in the nonseasonal model. This value will keep the number of infectives from falling to arbitrarily small values while not eliminating the divorce effect. We note that values of $I_b$ that are sufficiently large to eliminate the divorce effect would require $I_b$ to be roughly the same size as the endemic infection level.

It is well known that seasonally forced models are even more prone to having their numbers of infectives falling to low levels between outbreaks, with a background force of infection being commonly employed to counter this effect. Stronger seasonality magnifies this effect. Hence the background force of infection impacts the magnitude of the divorce effect, and given that the timing of control plays an important role in seasonal settings, there is an interaction between seasonality, the timing of the control, and the background force of infection in such cases. In general, as seasonality increases so does the difference between the maximum and minimum prevalence levels in the population. This results in an interaction between the background force of infection, the magnitude of seasonality, and the timing of the control determining the final magnitude of the divorce effect (Figures S8 and S23). This is important for predicting the magnitude of the divorce effect in real world situations, as there is a large amount of uncertainty associated with estimates of all three of these parameters. Importantly, below a specific background force of infection, the divorce effect is seen for all values of these parameters.

(a) (b)

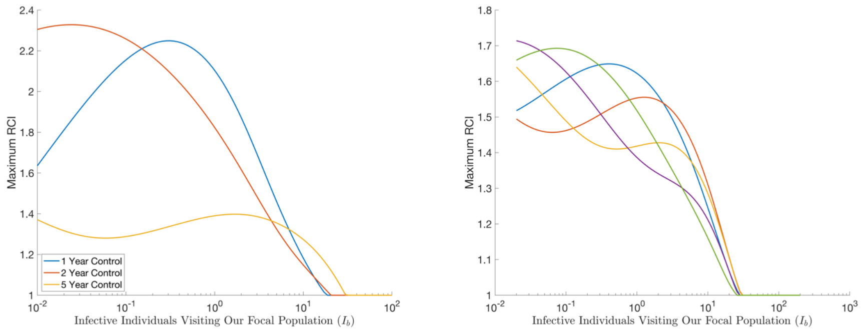


**Figure S23: Interaction between background force of infection, seasonality, timing of control, and the magnitude of the divorce effect.** (a) Increasing seasonality ($\beta_1 = .1$ in contrast to $\beta_1 = .02$ in Figure S22) increases the variation in the divorce effect seen at differing background force of infections. (b) Likewise, the timing of the start of the control (days 0, 73, 146, 219, 292) has a significant effect on the magnitude of the divorce effect seen at a particular background force of infection.

| | | |
|---|---|---|
| **Page 21: [1] Deleted** | **Brandon Hollingsworth** | **9/13/19 11:28:00 AM** |
| **Page 23: [2] Deleted** | **Brandon Hollingsworth** | **9/13/19 11:28:00 AM** |
| **Page 24: [3] Deleted** | **Brandon Hollingsworth** | **9/13/19 11:28:00 AM** |
